# Restoration of E-cadherin Expression Alters Metastatic Organotropism in Invasive Lobular Breast Carcinoma Models

**DOI:** 10.64898/2026.05.14.724680

**Authors:** Laura Savariau, Nilgun Tasdemir, Insa Thale, Ashuvinee Elangovan, Kai Ding, Dixcy Jaba Sheeba John Mary, Brent T. Schlegel, Jennifer M Atkinson, Jagmohan Hooda, Adrian V Lee, Steffi Oesterreich

## Abstract

Invasive lobular carcinoma (ILC) is the most frequently diagnosed special histological subtype of invasive breast cancer and accounts for 10 – 15% of all cases. The pathognomonic hallmark of ILC is the genetic loss of E-cadherin (*CDH1*) causing the disruption of adherens junctions and resulting in discohesive, linear growth. To better understand the role of E-cadherin in ILC metastasis, we generated three ILC cell lines, MDA-MB-134-VI, SUM44PE, and BCK4, with inducible E-cadherin expression, resulting in successful restoration of functional adherens junctions. E-cadherin expression reduced growth in 2D culture, and that effect was even greater in 3D ultra-low attachment (ULA) conditions where increased cell death was consistent with the previously described role of E-cadherin in anoikis. E-cadherin expression did not rescue the lack of migration and invasion of ILC cell line models; however, it decreased haptotaxis and increased adherence to Collagen I in SUM44 cells. There was no significant effect of E-cadherin expression on primary orthotopic tumor growth, but spontaneous metastasis to the reproductive tract, brain, and GI tract was reduced. Inhibition of metastasis to the reproductive tract and brain was also seen after tail vein injection of MDA-MB-134 E-cadherin-expressing cells. In summary, overexpression of functional E-cadherin in ILC models has some, but limited, effects on 2D growth *in vitro* and primary tumor growth *in vivo*, but there are pronounced effects on 3D ULA growth and metastases *in vivo*, with stronger effects on metastatic sites enriched in patients with ILC, especially the reproductive and GI tracts.

## Introduction

Breast cancer is a major public health concern with an estimated 319,750 new cases resulting in 42,680 deaths every year in the United States (*1*). In addition, breast cancer has the highest treatment costs of any cancer representing ∼14% of all cancer treatment costs in the United States (*2*). Despite tremendous advancement in breast cancer care with the discovery of endocrine therapies (*3*), 20-30% of patients treated with early stage breast cancer still develop metastatic disease (*4*).

Invasive lobular breast carcinoma (ILC) is the most frequently diagnosed special histological subtype of invasive breast cancer, and accounts for 10-15% of all breast cancer cases. There is an estimated 32,000 – 48,000 new patients diagnosed with ILC each year, and ranked separately, the disease would be the seventh most common cancer among women in the United States (*1*). ILC is characterized by a distinctive histology in which cancer cells grow in a single-file pattern, in stark contrast to breast cancer of “no special type” (NST, also known as invasive ductal carcinoma, IDC), where cohesive cells form bulky, palpable masses (*5*). This pattern of growth renders ILC difficult to detect and complicates surgical removal (*6*). In addition, ILC is more often multicentric and bilateral than NST, and displays unique pathognomonic features and sites of metastases. Multiple reports highlight the increased frequency of metastases to the ovaries, peritoneum, gastrointestinal tracts, and leptomeninges in patients with ILC compared to patients with NST (*7–9*), and our group also documented orbital metastases in patients with ILC (*10*). ILC are more frequently estrogen receptor (ER) positive than NST tumors, at rates of ∼90% for ILC and 60-70% for NST (*11–13*). Alongside a high rate of ER positivity, ILCs tumors tend to display HER-2 negativity and low Ki-67, which together are favorable initial prognostic and predictive factors. The majority of patients with ILC are treated with endocrine therapy, and although this shows high efficacy, long-term outcome is worse than those with ER+ NST due to higher rates of late recurrences (*14–16*). At the molecular level, other than the hallmark loss of *CDH1,* ILC tumors present with *PTEN* loss and increased PI3K/AKT signaling, loss of *TBX3* through mutations, and activating *FOXA1* mutations (*17*). Despite growing realization of unique features of lobular tumors, treatment guidelines are the same compared to patients with NST, highlighting the scarcity of clinical research dedicated to ILC.

The hallmark of ILC is the loss of *CDH1* mostly through truncating mutation, but also through mutations causing structural variants (*18, 19*) and in some cases *CDH1* methylation (*20*). E-cadherin is a transmembrane glycoprotein responsible for calcium-dependent cell adhesion. Its five extracellular cadherin repeats engage in trans-dimerization with cadherins on adjacent epithelial cells, while the intracellular domain interacts with α-, β-, and p120-catenins to anchor the cytoskeleton at adherens junctions (*21*). In many cancer types, E-cadherin loss is associated with epithelial-to-mesenchymal transition (EMT) and correlates with poor outcomes (*22*). On the contrary, E-cadherin has been shown to be required for viable cell dissemination and metastatic outgrowth in NST (*23*). In ILC, however, tumor cells retain an epithelial character despite *CDH1* loss, and EMT markers are largely absent (*24*). Here, we sought to define the role of E-cadherin in ILC progression, with particular focus on the unique metastatic organotropism characteristic of this disease. Using human ILC cell lines with doxycycline-inducible E-cadherin expression, we assessed the consequences of E-cadherin restoration on both *in vitro* phenotypes and *in vivo* metastatic behavior in xenograft models.

## Materials and Methods

### Cell culture

Cell lines were authenticated by the University of Arizona Genetics Core and mycoplasma tested in-house (Lonza #LT07–418). Following authentication lab stocks were generated which were used for this study. MDA-MB-134-VI (MDA-MB-134), MCF-7, T47D and MDA-MB-231 were obtained from the ATCC. SUM44PE (SUM44) was purchased from Asterand and BCK4 cells were developed as reported previously (*25*). Cell lines were maintained in the following media (Life Technologies) with 10% FBS: MCF7, T47D and MDA-MB-231 in DMEM, MDA-MB-134 in 1:1 DMEM:L-15, and BCK4 in MEM with nonessential amino acids (Life Technologies) and insulin (Sigma-Aldrich). SUM44 cells were maintained as described previously (*26*) in DMEM-F12 with 2% charcoal-stripped serum and supplements (Gemini Cat# 100-119). Cell lines were routinely tested to be *Mycoplasma* free. E-cadherin over-expression and E-cadherin knock out cells were generated as previously described (*27*). Estrogen and antiestrogen treatments were performed on hormone-deprived cells as described previously (*26*). Doxycycline (dox, Sigma-Aldrich) treatments were 1 µg/ml unless indicated otherwise.

### Cell sorting

To induce E-cadherin expression, cells were treated with dox for 24 hours before sorting. Cells were washed with PBS and detached with 2 mM EDTA before being spun down and washed in 3% BSA in PBS with 0.5 mM EDTA. The cells were incubated with Alexa Fluor 488 Mouse Anti-Human CD324 (E-cadherin) Clone 67A4 (BD Biosciences # 63570) for 45 min on ice and protected from light before pipetting through filtered tubes as required for sorting. The cells were sorted with FACSARIA Fusion sorter. Negative controls stained with Alexa Fluor 488 Mouse IgG1 K Isotype Control Clone (BD Biosciences #557782) were used for gating.

### Immunoblotting

Whole cell lysate samples for immunoblot analysis were collected using RIPA buffer (50mM Tris pH 7.4, 150mM NaCl, 1mM EDTA, 0.5% Nonidet P-40, 0.5% NaDeoxycholate, 0.1% SDS, 1× HALT cocktail [Thermo Fisher #78442]), then probe sonicated and centrifuged at 14,000 rpm at 4°C for 15 minutes. Protein concentrations were determined by BCA assay (Thermo Fisher #23225) and 40 µg of protein per sample was run on 10% SDS-PAGE gel along with PageRuler Plus Prestained Protein Ladder (Thermo Fisher Scientific, #26620). The gels were transferred onto PVDF membranes, which were then blocked in Odyssey PBS Blocking Buffer (LiCor #927–40000) for 1 hour at room temperature and then incubated in primary antibodies overnight at 4°C: E-cadherin (Cell Signaling Technology Cat#3195), α-catenin (Sigma-Aldrich Cat#C2081), β-catenin (Cell Signaling Technology Cat#9566), p120-catenin (BD Biosciences Cat# 610134), β-actin (Sigma-Aldrich Cat# A5441). Membranes were incubated in LiCor secondary antibodies for 1 hour at room temperature (anti-rabbit 800CW [LiCor #926–32211]; anti-mouse 680LT [LiCor #925–68020]; 1:10,000) and imaged with LiCOr Odyssey CLx Imaging system.

### Immunofluorescence confocal microscopy

Cells were seeded onto coverslips (Thermo Fisher Scientific #12-454-81), then treated with doxycycline for indicated amount of time. Coverslips were washed with PBS then fixed with ice-cold methanol for 15 minutes and washed again. Next, the coverslips were blocked in 5% BSA in 0.3% PBS-T for 1 hour, incubated with primary antibody diluted in blocking buffer overnight (Supplementary table 1_Antibodies), followed by Alexa Fluor secondary antibody incubation for 1 hour (anti-rabbit Alexa Fluor 488 [Life Technologies #A11070] and anti-mouse Alexa Fluor 546 [Life Technologies #A11018]; 1:200). Lastly, the coverslips were stained with Hoechst (Thermo Fisher Scientific #3570) diluted 1:10,000 in blocking buffer for 15 min, then mounted into slides with Prolong Diamond Antifade (Invitrogen P36961). Images were taken with Nikon A1, and images were processed with FIJI. E-cadherin colocalization with 120-catenin ratio was calculated with Nikon software (NIS-Elements AR).

### Immunohistochemistry (IHC) and Masson’s trichrome staining

Mouse organs were placed in cassettes overnight in 10% formalin (Sigma-Aldrich #HT501128), then switched to 70% ethanol until processing into wax blocks. Pitt Biospecimen Core assisted with paraffin-embedding and 5 µm sectioning onto slides. Slides were baked for at least 1 hour at 65°C, then deparaffinized and rehydrates by successive dipping of xylene (3×5min), 100% ethanol (3×5min), 95% ethanol (5min), 80% ethanol (3min), 70% ethanol (3min), PBS (3min), water (2×3min). Next, antigen retrieval was performed in Citrate buffer (sodium citrate–citric acid) in pressure cooker for 20 min. Slides were cooled before circling tissue with PAP pen (ab2601) followed by 5 min peroxidase activity blocking with hydrogen peroxidase (Abcam #ab64218). The slides were rinsed with PBS-T (0.5% Tween20), then incubated for 1 hour with blocking buffer made of 3% Bovine Serum Albumin (Sigma-Aldrich #A9647) in PBS-T. Next the slides were incubated overnight at 4°C with primary antibody diluted in blocking buffer. Anti-Cytokeratin 19 (Fisher Scientific #MS198P1), E-cadherin (CST #3195S), Ki-67 (Roche diagnostics #05278384001) were diluted 1:100. The next day, slides were rinsed 3 times in PBS-T before a 45min incubation with appropriate secondary antibodies (Mouse DAKO – K4001, Rabbit DAKO – K4011). Slides were next rinsed with PBS-T followed by staining with DAB (Agilent K3468) and counterstained in hematoxylin before rinsing in water and dehydration by dipping in 70% ethanol (3min), 95% ethanol (3min), 100% ethanol (3×3min) and lastly in xylene (3×3min). Slides were mounted with permount (Fisher Scientific #SP15100).

For trichrome staining the Trichrome, Masson, Aniline Blue Stain Kit (#9179A, Polysciences Inc.) was used according to manufacturer’s protocol, with minor adjustments. Paraffin-embedded tissue sections were first deparaffinized in xylene (3×3min), followed by rehydration through 100% and 95% ethanol (10 dips each). For mordanting, slides were incubated in preheated Bouin’s solution (50-60°C) for 1 hour and then cooled at room temperature for 5-10min. Slides were rinsed under running tap water and then distilled water. Nuclei were stained using freshly prepared Weigert’s iron hematoxylin for 10 min. The hematoxylin solution was prepared by mixing acidified chloride (Solution B) with 1% alcoholic hematoxylin (Solution C). Following nuclear staining, slides were washed well under running tap water and rinsed with distilled water. Cytoplasm and muscle fibers were stained with Biebrich Scarlet-Acid Fuchsin Stain (Solution D) for 2 min. After rinsing in distilled water, sections were differentiated in phosphomolybdic-phosphotungstic acid solution (Solution E) for 10-15 min, then transferred directly to Aniline Blue (Solution F) for 5 min to stain collagen. Slides were rinsed with distilled water, treated with 0.5% aqueous acetic acid (Solution G) for 3-5 min, dehydrated through 95% (2×10 dips) and 100% (2×10 dips), cleared in xylene (3×10 dips). Slides were mounted with permount (Fisher Scientific #SP15100). Images were taken with the Nikon Eclipse 50i and images were analyzed using ImageJ.

### E-cadherin adhesion assay

Recombinant Human E-cadherin Protein, CF (R&D Systems #8505-EC) was reconstituted in Coating Buffer (PBS with calcium and magnesium (Corning #21-030-CV)) (250µg/ml). Non-treated 96 well plates (Fisher #12-565-226) were used for a 2-fold serial dilution of E-Cadherin coating with a starting coating concentration of 6μg/ml. For background control, Coating Buffer only wells were used. E-cadherin coated plates were incubated overnight at 4°C, then blocked with Blocking Buffer (10mg/ml BSA and PBS with calcium and magnesium (Corning #21-030-CV)) for 1 hour at 37°C. Dox pretreated cells were disassociated with 2mM EDTA, washed with blocking buffer, stained with 1μM Calcein-AM (Thermo Fisher # C1430) for 30 min at 37°C, then seeded at 5×10^5^ cells/ml for MCF7 and 1×10^6^ cells/ml for ILC cell lines. Following a 12 hour incubation, the plates were read before and after 3 PBS wash with Perkin-Elmer Victor X4. The average percent adhesion was calculated (OD after wash)/(OD before wash) *100.

### Apoptosis assay and cell cycle assay

Cells were seeded at 300,000/well in 6-well 2D and ULA plates in triplicates. After 4 days, for apoptosis assessment cells were stained with APC-Annexin V (BD Biosciences; #550474) and PI in 1X Annexin binding buffer for 15 min at room temperature. For cell cycle analysis, cells were stained with Hoechst (Thermo Fisher Scientific) at 20 mg/ml for 30 min at 37°C and then briefly with Propidium Iodide (BD Biosciences) to gate on viable cell. Samples were acquired on an LSR II Flow cytometer (BD Biosciences) and analyzed using BD FACSDiva and FlowJo software (BD Biosciences).

### Transwell migration, haptotaxis and invasion assays

Cells were pre-treated with dox for 24 hours, serum starved overnight, detached with 2mM EDTA and 300,000 cells were plated into each 8-µm inserts (Thermo Fisher Scientific and Cell Biolabs), Quantification with crystal violet was performed following manufacturer’s protocol. Images of inserts were taken on an Olympus SZX16 dissecting microscope. For migration and invasion assays, 10% FBS was added to the bottom chambers. No chemoattractant was added for haptotaxis to Collagen I.

### Growth assay

2D and ULA growth assays were performed as previously described (*28*). ILC (15,000/96-well; 300,000/6-well) and IDC (5,000/96-well; 100,000/6-well) cells were seeded in cell culture treated (Thermo Fisher Scientific) or ULA (Corning Life Sciences) 96-well plates. Cells were assayed using CellTiter-Glo (Promega) or FluoReporter Blue Fluorometric dsDNA Quantitation Kit (Invitrogen). Data was captured on a Promega GloMax or Perkin Elmer Victor X4 plate reader.

### Estrogen response

Estrogen induced growth experiments were performed following estrogen deprivation with phenol-red free iMEM with 5 or 10% CSS, washing 3 times a day for 3 days. The cells were plated at 5,000 cells/well for IDCs and 15,000 cells/well for ILCs and stimulated with 1nM estradiol (Sigma Aldrich, #E8875) or vehicle (ethanol). The plates were read with CellTiter-Glo (Promega, #PR-G7573) or PrestoBlue (Thermo Fisher Scientific, #A13262) at day 1 and 8 to calculate fold change. For qPCR, cells were deprived in the same manner described above and then treated for 4hours with 1 nM E2 or vehicle. RNA extraction was conducted following manufacturer protocol for RNeasy Mini Kit (Qiagen #74106). RT was conducted following the manufacturer protocol for PrimeScript RT Master Mix (Takara Bio, RR036B) and qPCR was performed in 396 well plates using 50ng of RNA per sample in triplicate with the SsoAdvanced Universal SYBR SMX (Bio-Rad, # 1725275) and run in the CFX384 PCR Detection System (Bio-Rad).

### Animal Experiments

All experiments were conducted on 4-week-old female NOD.Cg-*Prkdc^scid^ Il2rg^tm1Wjl^*/SzJ (NSG) mice purchased from the Jackson Laboratory (RRID:IMSR_JAX:005557) under the University of Pittsburgh IACUC protocol #20226775. All mice were implanted subcutaneously on the back of the neck with a 17β-estradiol (0.5mg, 90 days release) from Innovative Research of America the day before cell injections. The pellet implant procedure was conducted under sterile conditions with the mice under isoflurane anesthetic (Patterson Veterinary, #78938441). Subcutaneous injection of Ethiqa XR (50ul of 3.25mg/kg) and Ketofen (100ul of 5ug/ml) (covetrus #072117, #005487) were given for pain management as needed. After the 90 days, the mice were supplied water containing 8 µg/ml of estradiol. All mice were supplied doxycycline (dox) diets (Envigo TD.130141) starting the day of the 17β-estradiol pellet implantation.

Mammary fat pad injections (MFP) were performed in the 4^th^ fat pad, with 3 million cells mixed 1:1 in media and Matrigel (Corning, #356237) using a 26G syringe. The cells were treated with dox for 24 hours before all the injections. Tumor volumes were monitored by caliper measurements weekly. A first cohort of mice with n=10 for each group (EV and CDH1) for both MDA-MB-134 and SUM44 underwent MFP injection to monitor the effect of E-cadherin on tumor growth and testing E-cadherin level expression *in-vivo*. For this experiment the cells were stably labelled with RFP-Luciferase by infection with the pLEX-TRC210/L2N-TurboRFP-c lentivirus plasmid (*29*). Upon the tumors reaching 1000 mm^3^ or the mice showing sign of advanced disease, mice were euthanized.

A second cohort of mice with n=10 for each group (EV and CDH1) for both MDA-MB-134 and SUM44 received tail vein (TV) injections. For this experiment the cells were stably labelled with RFP-Luciferase by infection with the pLEX-TRC210/L2N-TurboRFP-c lentivirus plasmid (*29*) which allowed tracking of the metastatic dissemination using IVIS200 *in vivo* imaging system (PerkinElmer, #124262) following intraperitoneal injection of D-luciferin (Gold Bio) (100 ul, 150 mg/kg) and bioluminescence was measured. Cells were exposed with dox for 24 hours prior to injection. Mice were placed in a restraining device which allowed easy access to the tail. A heat lamp was used to dilate the vein, and approximately 2 million cells in 100ul of PBS were injected using a 26G needle. The mice were monitored weekly using IVIS imaging. For MDA-MB-134 and SUM44, mice were sacrificed ∼11 and ∼14 weeks post TV injection, when the first mice of the cohort required euthanization as per approved IACUC protocol.

Another cohort of mice with n=12 for each group (EV and CDH1) for both MDA-MB-134 and SUM44 underwent MFP injection with the above-mentioned RFP-Luciferase cells followed by tumor removal when the tumor reached 200-300 mm^3^ to test the effect of E-cadherin on metastatic dissemination. Mice were placed under anesthesia for the duration of the primary tumor removal surgery, and the above mentioned Ketofen and Ethiqa XR were given for pain management. The surgery was conducted under sterile conditions; the fat pad was shaved and cleaned with povidone-iodine and 70% ethanol. The wound was clipped and stitched if necessary. Mice were closely monitored following surgery, and the clips were removed 10-14 days post-surgery and were euthanized at approximately ∼40 ± 8 days, and ∼60 ± 13 days following surgery for SUM44 and MDA-MB-134 respectively. The end point was determined after the first mouse of each cohort underwent surgery and showed signs of advanced disease that required euthanasia. Endpoints were kept as similar as possible in each cell line after surgery for a fair comparison of the metastatic spread.

At the designated endpoint, each animal was imaged, and relevant organs were harvested. Excised organs containing metastatic lesions were placed in a 24-well plate for ex vivo fluorescence imaging. Immunofluorescence (IF) was performed to detect the RFP signal, using the IVIS Lumina X5 imaging system equipped with appropriate filters for RFP (excitation/emission: ∼558/583 nm).

### Survival and bar plots and statistical tests

GraphPad was used to make the survival plots, and log-rank Mantel-Cox test was performed to assess significance. GraphPad was also used for the bar plots representing the incidence of metastatic lesions. Incidence of metastasis per organs comparing EV vs CDH1 was reported as categorical (yes or no) and thus Fisher’s exact test in GraphPad was used to test statistical significance at each organ site. Statistical significance for the growth assays, apoptosis and haptotaxis experiments were assessed using two-way ANOVA test, t-test and unpaired t-test respectively.

Metastatic incidence was compared between CDH1_OE and EV groups using Fisher’s Exact Test, with each organ site per mouse treated as a binary outcome (yes/no metastasis per organ, per mouse). Raw counts were pooled across five organ sites (Spleen, GI, Kidneys, Brain, and Reproductive Tracts) per cohort, giving a total of N_mice × 5 observations per group. Fisher’s Exact Test was applied independently to each of the four experimental cohorts (SUM44 TV, MM134 TV, SUM44 MFP, MM134 MFP) using the base R fisher.test() function, which computes exact p-values via the hypergeometric distribution without large-sample assumptions. Odds ratios and 95% confidence intervals were extracted directly from the test output. Benjamini-Hochberg correction was applied across the four comparisons using p.adjust(method = “BH”) to control the false discovery rate. Data cleaning, statistical testing, and visualization were performed using the tidyverse (v2.0.0) and ggplot2 (v4.0.2) packages. Statistical analyses were performed in R version 4.5.0.

## Results

### Doxycycline induced E-cadherin expression in ILC restores adherens junctions

To study the role of E-cadherin loss in the context of ILC, we generated three human ILC cell lines (MDA-MB-134, SUM44 and BCK4) with inducible E-cadherin overexpression. Immunoblotting of the 3 cell lines after 24 hours dox treatment shows successful induction of E-cadherin expression (Figure 1A). We further observed increased expression of p120-catenin, α-catenin, and β-catenin upon CDH1 induction, suggesting that E-cadherin expression alone is sufficient to restore adherens junction formation. E-cadherin (pink) and p120-catenin (green) co-localized upon exposure to dox suggesting the successful restoration of functional adherens junctions. Immunofluorescence imaging revealed that dox-induced E-cadherin expression drove a rapid transition from discohesive single cells to cohesive clusters within 24 hours in MDA-MB-134 and SUM44 cells, consistent with the restoration of adherens junctions (Figure 1C). In contrast, BCK4 cells required up to 86 hours of treatment before this phenotypic transition was observed (Figure 1B). Quantification of colocalization of p120-catenin with E-cadherin (Supplementary Figure S1A, B and C) and immunoblotting of E-cadherin and adherens junction proteins (Supplementary Figure S1D, E, and F) suggest that 24 hours of dox treatment is necessary for E-cadherin to reach the plasma membrane and induce adherens junction formation in MDA-MB-134 and SUM44. A defining function of adherens junctions is to mediate intercellular adhesion through trans-engagement of extracellular cadherin repeats on neighboring cells. Consistent with this, MDA-MB-134-CDH1 and SUM44-CDH1 cells pre-treated with dox for 24 hours adhered to E-cadherin-coated plates nearly as efficiently as the E-cadherin-positive control line MCF7 (Figure 1D), further validating the functional restoration of adherens junctions in our models.

**Figure 1.**
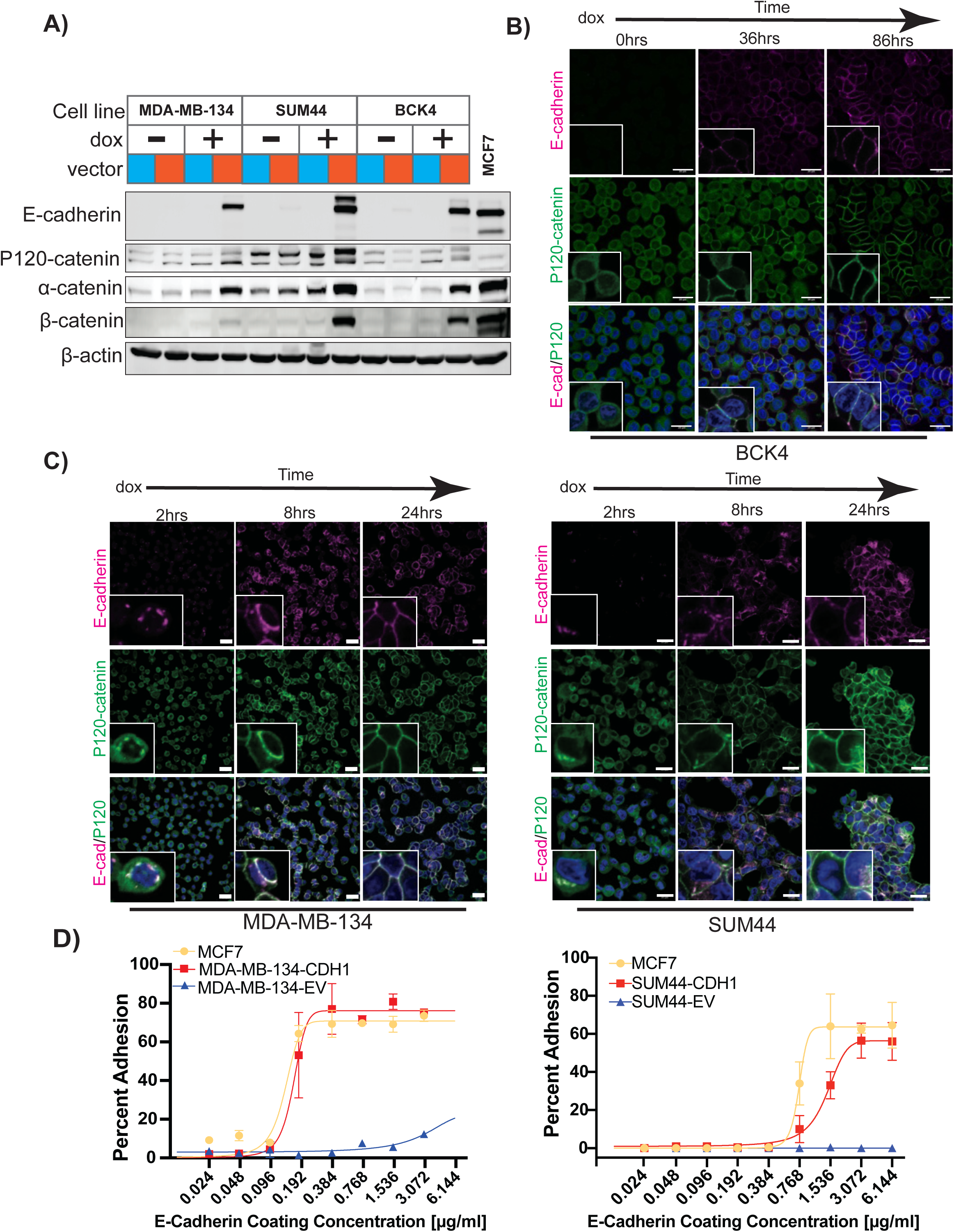
Doxycycline induced E-cadherin expression restores adherens junctions in ILC cell lines. A) Immunoblot using antibodies for E-cadherin, α, β and p120-catenins on whole cell lysate of indicated cells. EV and CDH1 cells are represented in blue and red, respectively, and with or without 24 hours dox treatment. B) & C) Immunofluorescence confocal microscopy of p120-catenin (green) and E-cadherin (pink) of pINDUCER20-CDH1 indicated cell lines with increasing duration of dox treatment. Dapi (blue) was used for nuclei staining. Scale bars represent 20 μm. D) E-cadherin adhesion assay with indicated cells lines. Cells were pre-treated with dox for 24 hours prior to the assay.

### E-cadherin expression reduces anoikis resistance and haptotaxis, and increases adhesion

Prior studies have demonstrated that E-cadherin loss re-localizes p120-catenin to the cytoplasm and nucleus, thereby enabling anoikis resistance (*30, 31*). We have recently shown that a panel of ILC cell lines exhibits pronounced anoikis resistance, proliferating robustly under ULA conditions that force cells to grow in suspension (*28*). We therefore hypothesized that restoring E-cadherin expression would reverse this anoikis-resistant phenotype. In MDA-MB-134, E-cadherin overexpression significantly inhibited 2D growth, and totally blocked growth in ULA (Figure 2A). Similar effects were seen in SUM44 cells although the effect size was smaller. Further analysis of the reduced ULA growth in MDA-MB-134 showed that E-cadherin expression induced cell death (Figure 2B and Supplementary Figure S2A). While ULA conditions alone increased apoptosis relative to 2D, E-cadherin re-expression further and specifically amplified both early and late apoptotic populations, demonstrating that E-cadherin sensitizes MDA-MB-134 cells to detachment-induced cell death.

**Figure 2.**
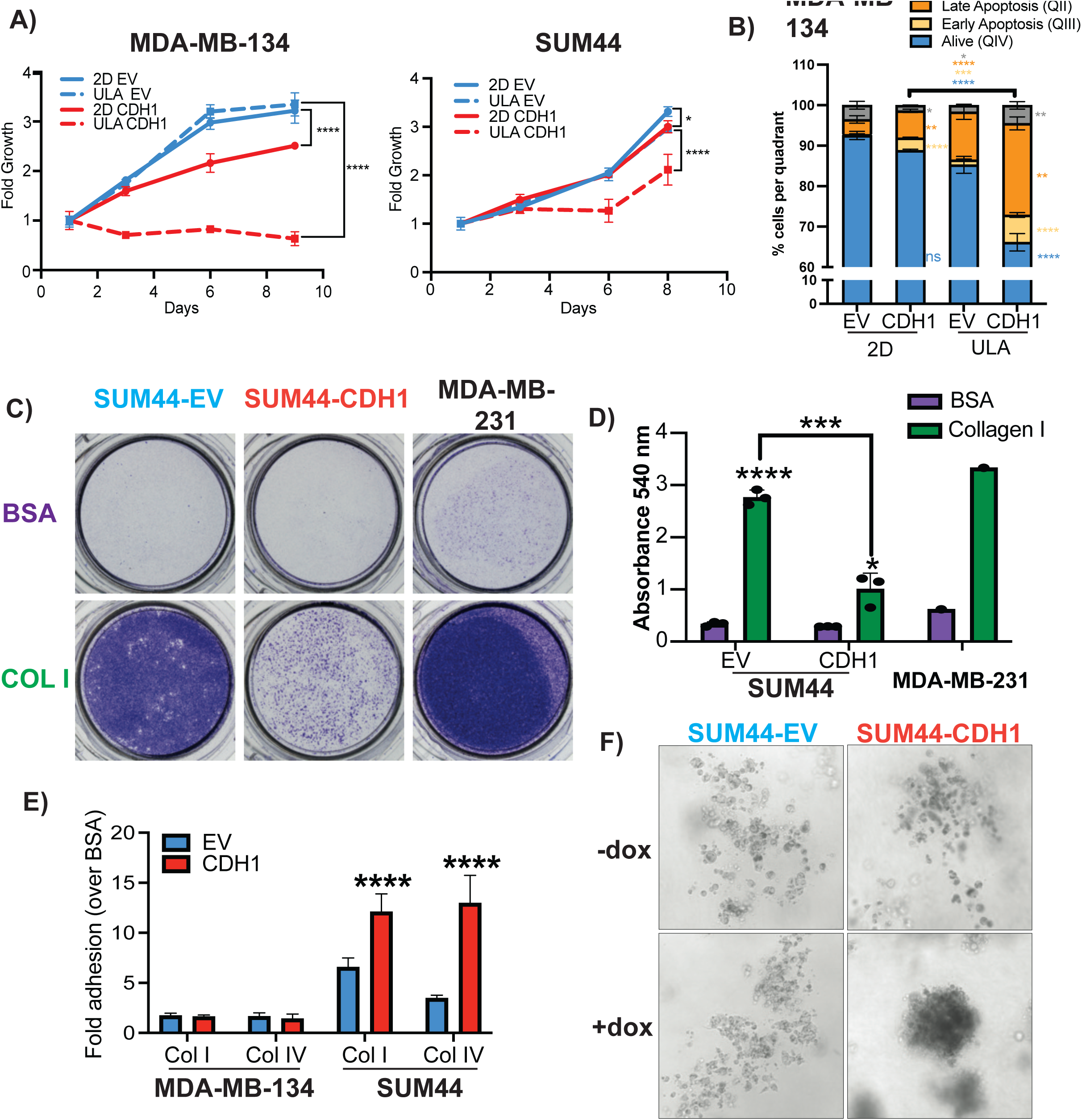
Phenotypic alteration with E-cadherin expression in ILC. A) Growth in 2D (line solid) and ULA (line dashed) culture of EV (blue) and CDH1 (red) indicated cell lines. Graphs show representative data of at least 2 experiments (n=6). P-values are from two-way ANOVA comparison between EV and CDH1 for each condition. *, P ≤ 0.05; **** P ≤ 0.0001. B) Quantification of the viable (Q4:Annexin V-/PI-) population of MDA-MB-134 EV/CDH1 in 2D and ULA. Data display mean percentage +/-standard deviation to the 2D condition. Graphs show representative data from two experiments (n=3). p-values are from t-test. *, P ≤ 0.05; **, P ≤ 0.01, ***, P ≤ 0.001; ****, P ≤ 0.0001. Statistical results next to the bars indicate comparison of EV vs CDH1 in each 2D and ULA condition, and statistics above the bars indicate comparison of CDH1 groups in 2D vs ULA. C) Images of crystal violet stained inserts with SUM44-EV and CDH1 cells from haptotaxis assay toward Collagen I for 72 hours. D) Quantification of SUM44 haptotaxis assay showing representative data from 4 experiments (n = 3). P values are from unpaired t-test. *, P ≤ 0.05; ****, P ≤ 0.001; ****, P ≤ 0.0001. E) SUM44 but not MDA-MB-134 show increased adherence to Collagen I (Col I) and Collagen IV (Col IV). Representative data from 3 experiments (n=6). P values are from unpaired t-test. ****, P ≤ 0.0001.F) SUM44-CDH1 with dox form tight sphere compared to -dox.

Dox treatment consistently altered proliferation of BCK4 EV control cells (Supplementary Figure S2B), limiting reliable interpretation of E-cadherin-specific effects; consequently, the BCK4 model was excluded from further phenotypic analyses.

We next evaluated migration, invasion, and adhesion using a panel of *in vitro* assays. ILC cell lines show limited migratory and invasive capacity *in vitro* (*28*), and E-cadherin re-expression did not rescue these phenotypes (Supplementary Figure S3A–D). However, E-cadherin re-expression did affect haptotaxis in SUM44 cells, a form of directional migration driven by gradients of substrate-bound adhesion ligands or chemoattractants. To assess haptotaxis, transwell inserts were coated on their undersurface with Collagen I and placed in wells containing growth-factor-free media, permitting only cells in direct contact with Collagen I to migrate through. SUM44 cells exhibited robust haptotaxis toward Collagen I, which was significantly diminished by E-cadherin expression (Figure 2C&D). To determine whether direct substrate contact was required, cells were seeded on uncoated transwell inserts placed above Collagen-I-coated plates (Supplementary Figure S4A). Neither SUM44 EV nor E-cadherin-expressing cells migrated through the inserts under these conditions (Supplementary Figure S4B&C), confirming that direct contact with Collagen I is necessary to initiate haptotaxis. Haptotaxis toward fibronectin was also reduced by E-cadherin re-expression in SUM44, though this did not reach statistical significance (Supplementary Figure S5A–C). These findings are consistent with our prior observation that E-cadherin loss in the NST cell lines MCF7 and T47D increased haptotaxis to Collagen I (*32*). Although the effect was not statistically significant, E-cadherin expression appeared to promote haptotaxis toward fibronectin, in addition to Collagen I (Supplementary Figure S5D&E).

In adhesion assays, E-cadherin overexpressing SUM44 cells demonstrated significantly greater adherence to both Collagen I and Collagen IV compared to SUM44 control cells (Figure 2E). When SUM44-CDH1 cells were embedded in high-density Collagen I, they formed compact spheres, in contrast to the discohesive growth pattern of the control cells (Figure 2F).

### Lack of significant effects of E-cadherin re-expression in ILC cells on *in vivo* primary tumor growth

To assess the effect of E-cadherin re-expression on *in vivo* tumor growth, we stably labeled the dox-inducible cell lines with RFP and Luciferase to enable real-time tracking by IVIS bioluminescence imaging. Labeling had no effect on E-cadherin expression or proliferation (Supplementary Figure S6A&B). The labeled MDA-MB-134 (EV RFP/Luc and CDH1 RFP/Luc) and SUM44 (EV RFP/Luc and CDH1 RFP/Luc) cells were injected into NSG mice (Figure 3A, left panel).

**Figure 3.**
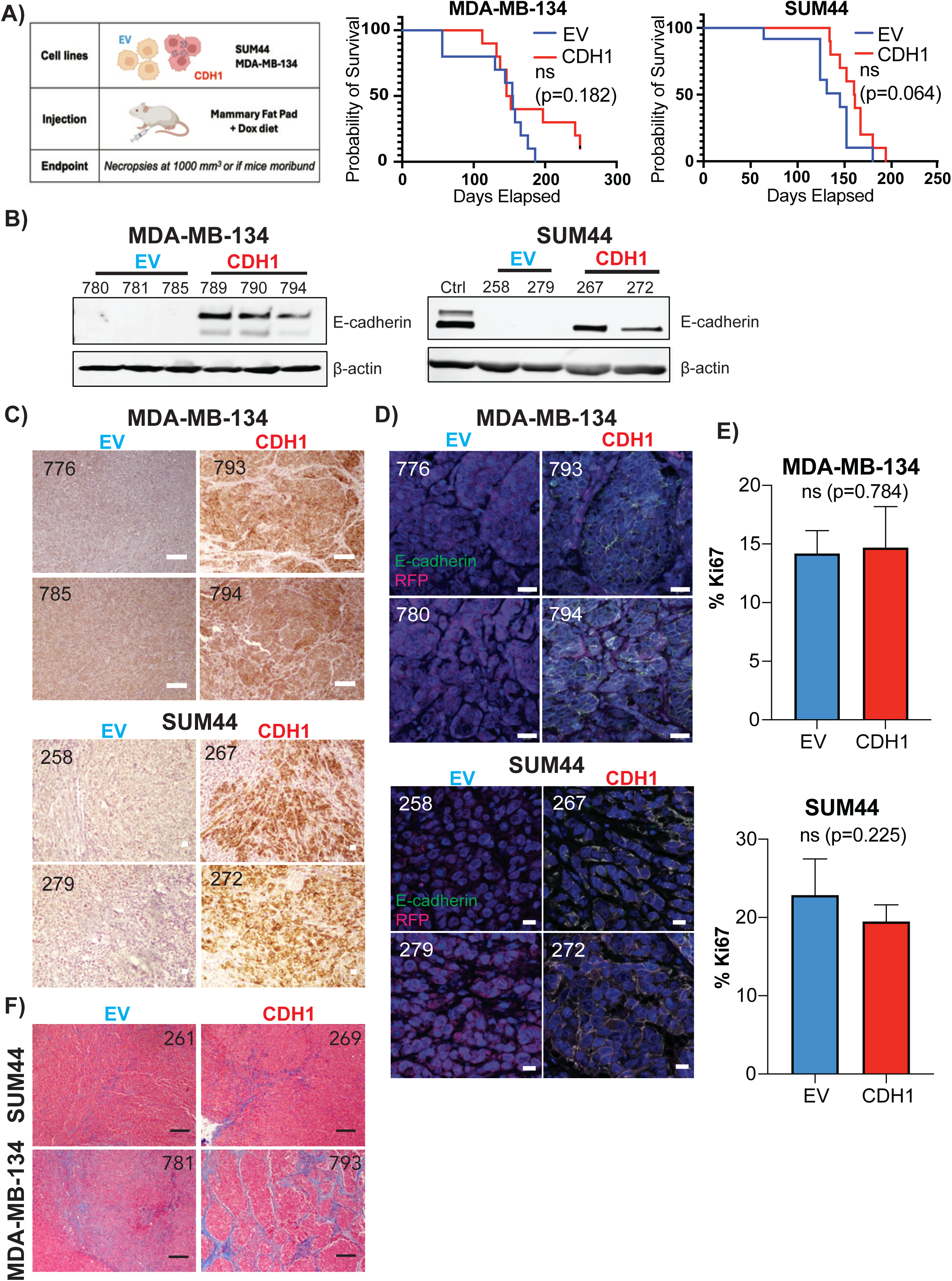
E-cadherin overexpression in ILC cells does not affect *in vivo* primary tumor growth, but alters morphology. A) Schematic of first mice experiment where NSG mice were placed on dox diet and E2 pellets were subcutaneously placed before mammary fat pad injection of EV and CDH1 cells with both SUM44 and MDA-MB-134 cell lines (n=10 for each group). Survival is defined as tumor reaching 1000 mm^3^ or mice succumbing to disease. B) Immunoblotting for E-cadherin from primary tumor of 2/3 mice from each EV and CDH1 group. Control is lysate from SUM44-CDH1 cell line. C) Immunohistochemistry for E-cadherin in primary tumor of 2 mice from each EV and CDH1 group. Scale bars represent 10µm. D) Immunofluorescence microscopy from FFPE fixed tumors for E-cadherin in green and RFP in pink. Scale bars represent 20µm. E) Quantification of nuclear Ki67 detected in 5 tumors. Quantification done with ImageJ and p-values are from unpaired t-test. ns; not significant. F) Representative Masson’s trichome staining (collagen in blue) of SUM44 and MDA-MB-134 tumors from each EV and CDH1 group. Scale bars represent 100 µm.

Primary tumor growth was not significantly affected by E-cadherin re-expression, though a non-significant trend toward reduced growth was observed for both MDA-MB-134 (p=0.182) and SUM44 (p=0.064) (Figure 3A). Expression of E-cadherin was confirmed by immunoblotting and IHC (Figure 3B&C, Supplementary Figure S7A&B). IHC revealed that E-cadherin expression in MDA-MB-134 tumors induced distinct histological changes, with formation of larger, more irregularly shaped cell clusters compared to the uniform, diffuse architecture of EV control tumors. In SUM44 tumors, the effect was more subtle; however, immunofluorescence staining revealed focal clustering of E-cadherin-positive cells in discrete regions, suggesting partial reorganization of cell–cell contacts *in vivo* (Figures 3C&D). Consistent with the lack of significant effects on primary tumor growth, Ki67 levels did not differ significantly between EV and E-cadherin-expressing tumors (Figure 3E, Supplementary Figure S7C&D).

To evaluate extracellular matrix (ECM) remodeling, Masson’s trichrome staining was performed on MFP tumors from EV and E-cadherin-expressing SUM44 and MDA-MB-134 cells (Figure 3F, Supplementary Figure S8A&B). Collagen content was quantified as the percentage of stained area (Supplementary Figure S8C&D). E-cadherin re-expression was associated with a trend toward more intense and widespread collagen deposition, though this did not reach statistical significance. In MDA-MB-134 tumors, E-cadherin-expressing samples additionally displayed collagen delineating defined cellular structures, a feature absent in EV controls.

### Tail vein injection of E-cadherin re-expressing ILC cells shows reduces metastasis to the reproductive tract

To examine the effect of E-cadherin on metastases, we injected RFP/Luc labelled MDA-MB-134 and SUM44 EV and E-cadherin expressing cells via tail vein (TV) route, monitored weekly by IVIS imaging to track disease burden, and then sacrificed the mice at the same time (Figure 4A). Metastatic incidence was assessed by RFP signal detection across the reproductive tract, brain, kidneys, GI tract, spleen, liver, and lungs.

**Figure 4.**
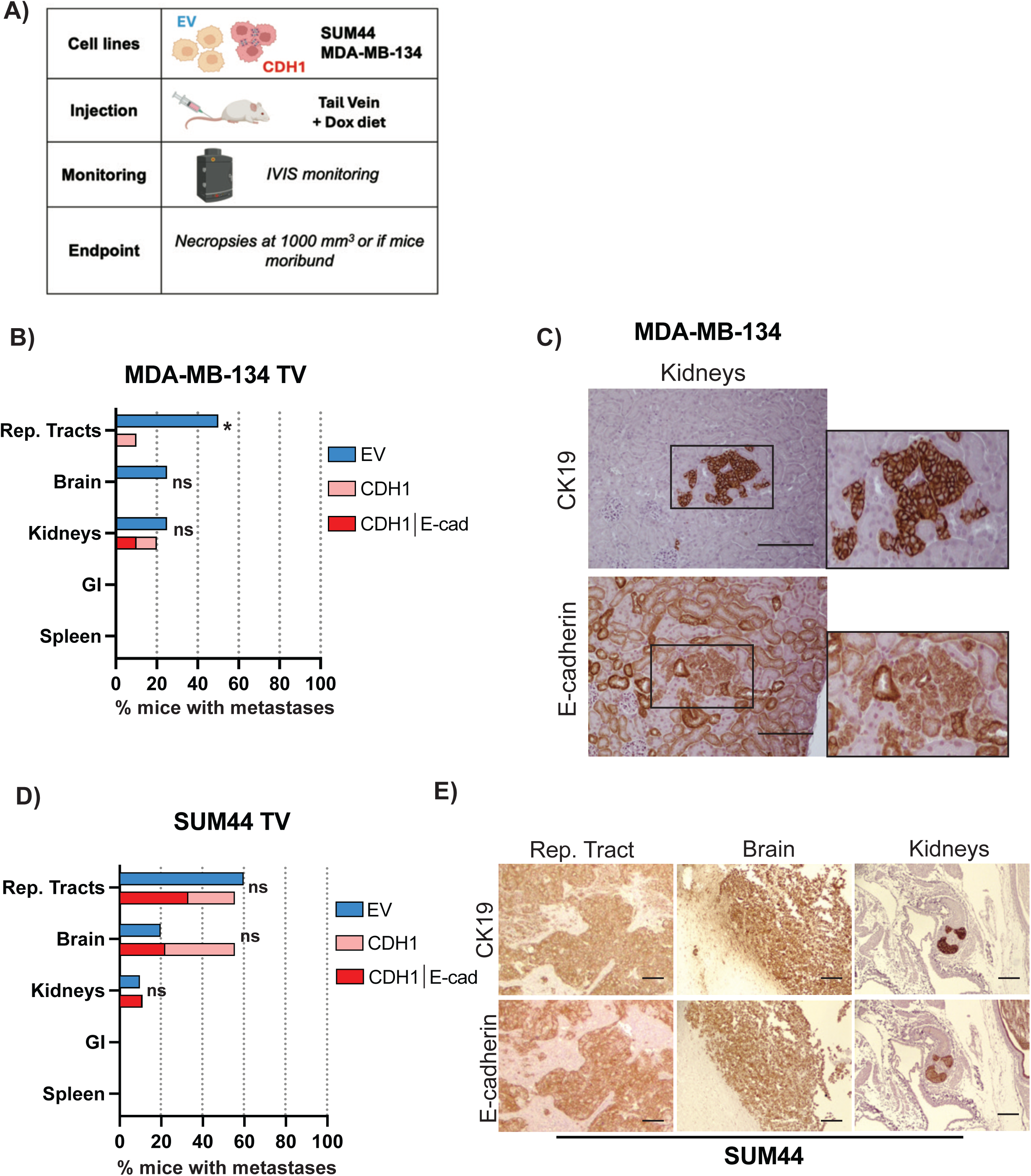
E-cadherin expression reduces metastatic seeding and outgrowth. A) Schematic of mice experiment. EV and CDH1 cells from both MDA-MB-134 and SUM44 were injected in the tail vein of mice which were then monitored by IVIS imaging weekly until harvest. B) & D) Incidence of tumors per organs for mice injected with EV cells in blue and for mice injected with CDH1 cells in light red. Dark red represents tumors in which E-cadherin expression was confirmed. C) & E) IHC staining for human marker CK19 and E-cadherin for tumors from mice injected with CDH1 cells for indicated organs for both cell lines MDA-MB-134 and SUM44. Scale bars represent 100 µm. Statistical analysis was performed using Fisher’s exact test (two-sided). *, P ≤ 0.05; ns, not significant.

At time of sacrifice (∼11 weeks post-injection), MDA-MB-134 EV cells (N=8) showed metastases in the reproductive tract (ovaries and uterine horn), brain, and kidney (Figures 4B). In contrast, mice injected with E-cadherin expressing MDA-MB-134 cells showed reduced metastases to the reproductive tract and brain (Figure 4B). RFP-positive organs were stained with anti-E-cadherin and the human-specific marker anti-CK19 antibodies to confirm and localize metastatic lesions. All EV metastases were E-cadherin-negative, as expected given their ILC origin. Notably, many metastases arising from injection of E-cadherin expressing cells (Figure 4C) were E-cadherin-negative, implying selective pressure favoring the outgrowth of E-cadherin-negative variants (Supplementary Figure S9A&B). For example, of the two kidney lesions observed in this group, only one stained positive for E-cadherin (Figure 4B&C). We did not observed any lung metastases, similar to what we had recently reported (*33*). Assessment of liver metastases was confounded by two factors: their presentation as microlesions, and the high endogenous E-cadherin expression in normal mouse liver, which precluded reliable IHC confirmation.

There was no significant effect of E-cadherin overexpression on metastases in the SUM44 model (Figure 4D&E). Of note, just like in the MDA-MB-134 model, some metastases from the SUM44 CDH1 TV injection were found to be E-cadherin negative (Supplementary Figure S9B). A trend toward reduced reproductive tract metastasis was observed in mice with confirmed E-cadherin overexpression in SUM44 cells. Overall, when analyzing total metastatic incidence across all organs, E-cadherin overexpression significantly reduced metastasis in the MDA-MB-134 TV model (OR=0.081, p=1.82×10^−2^, BH-adjusted Fisher’s exact test), but not in the SUM44 TV model (OR=0.703, p=6.52×10^−1^).

### E-cadherin expression alters organotropism following MFP injection

We next tested effects of E-cadherin on metastases after injection of EV and E-cadherin expressing ILC cells into the mammary fat pad (MFP) (Figure 5A, and Supplementary Figure S10A). The overall metastatic pattern following MFP injection recapitulated that of the TV injection for both MDA-MB-134 (Figure 5B&C) and SUM44 (Figure 5D&E), with predominant involvement of the reproductive tract, brain, and kidneys; the MFP model additionally yielded lesions in the GI tract and spleen. Overall, in this experiment of injection of cells directly into the MFP, more tumors from the CDH1-group lacked E-cadherin overexpression (Supplementary Figure S10 B&C).

**Figure 5.**
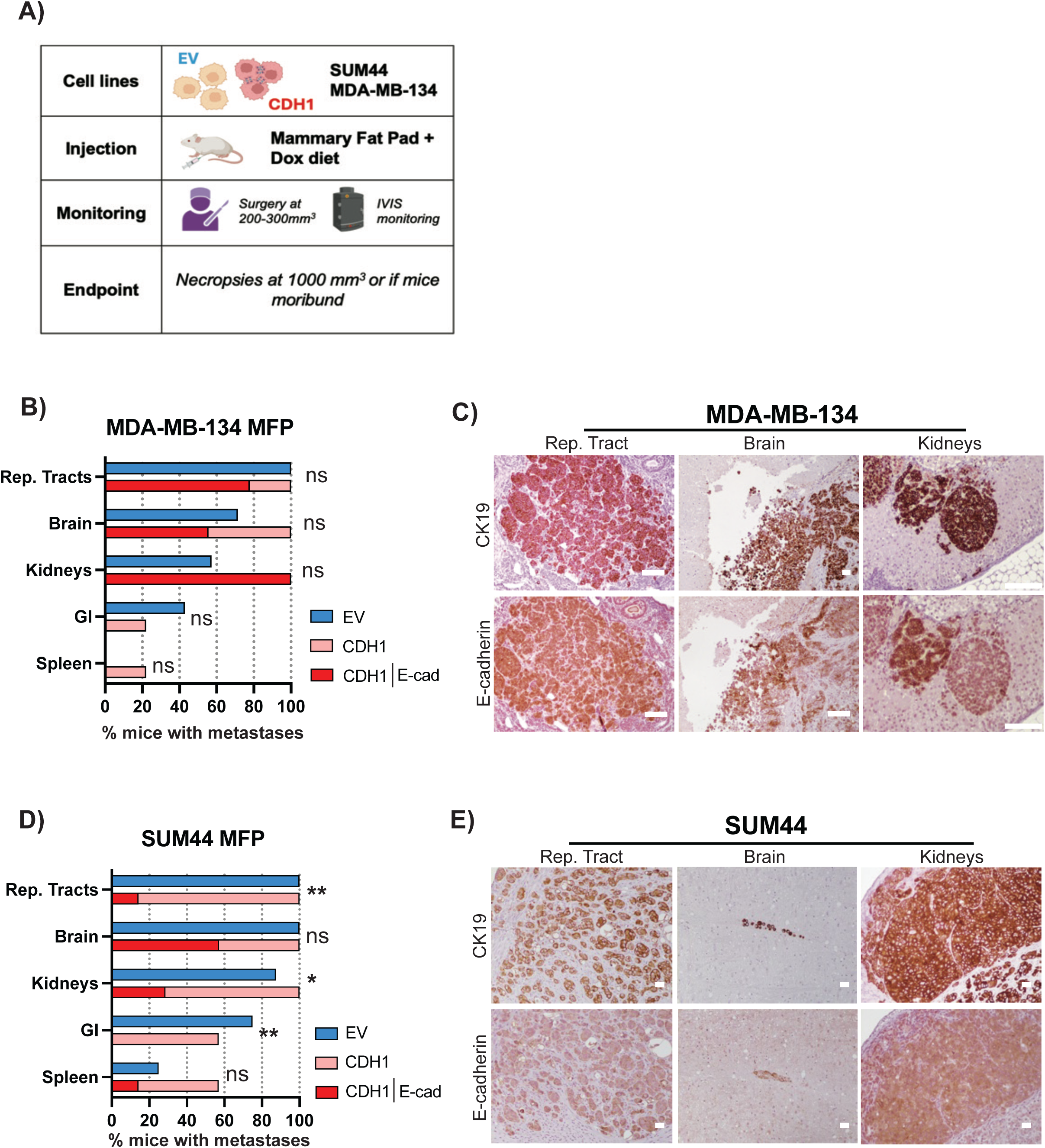
E-cadherin expression reduces metastatic dissemination and alters organotropism. A) Schematic of experimental settings B) & D) Incidence of tumors per organs in blue for mice injected with EV cells, light red for mice injected with CDH1 cells and in dark red incidence of tumors in which E-cadherin expression was confirmed. C) & E) IHC staining for human specific CK19 and E-cadherin of tumors from mice injected with CDH1 cells for indicated organs for both cell lines MDA-MB-134 and SUM44. Statistical analysis was performed using Fisher’s exact test (two-sided). *, P ≤ 0.05; **, P ≤ 0.01; ns, not significant.

In the MDA-MB-134 cohort, no significant differences in metastatic incidence were observed between EV (N=7) and E-cadherin-overexpressing (N=9) groups. Trends toward reduced metastasis to the brain, reproductive tract, and GI tract were present but did not reach significance, and E-cadherin re-expression was associated with a non-significant increase in kidney metastatic incidence.

In contrast, E-cadherin overexpression significantly altered the metastatic pattern in the SUM44 model (EV: N=8; CDH1: N=7). Metastatic incidence was significantly reduced in the reproductive tract and kidneys, with a complete absence of GI tract metastases in mice injected with E-cadherin expressing SUM44 cells. Non-significant trends toward reduced metastasis to the brain and spleen were also observed.

Statistical analysis of total metastatic incidence revealed an approximately 11-fold reduction in the odds of metastasis in SUM44 CDH1-expressing mice compared to EV controls (OR=0.090, p=1.03×10⁻⁵, BH-adjusted), while no such effect was detected in the MDA-MB-134 MFP cohort (19/35 vs 21/45; OR=1.352, p=6.52×10⁻¹).

## Discussion

The role of E-cadherin in breast cancer progression remains an area of active investigation. While the prevailing view holds that E-cadherin loss promotes EMT and thereby facilitates metastasis (*34*), ILC, a cancer defined by *CDH1* loss, paradoxically exhibits a propensity for late recurrence. As demonstrated by Padmanaban et al., NST cells retain E-cadherin, which is indispensable for critical steps of the metastatic cascade (*23*), and conversely, its loss impairs metastatic spread in that context. These contrasting observations underscore the context-dependence of E-cadherin’s role in cancer progression and motivated us to investigate its function specifically in ILC.

Using dox-inducible E-cadherin expression in human ILC cell lines, we showed that restoration of E-cadherin resulted in functional adherens junctions. E-cadherin re-expression suppressed both 2D and ULA growth, with a markedly stronger effect under ULA conditions, mirroring the differential growth behavior of NST cell lines (*28*). The disproportionate growth suppression under ULA conditions, together with the induction of apoptosis in E-cadherin-expressing MDA-MB-134 cells, confirmed the critical role of *CDH1* loss in ILC anoikis resistance. Restoration of E-cadherin re-sequestered p120-catenin to the plasma membrane, causing a number of downstream effects including the previously described relief of Kaiso-mediated transcriptional repression, thereby reversing anoikis resistance (*35*). Given that E-cadherin expression substantially impaired growth in both 2D and ULA conditions, particularly in MDA-MB-134, the absence of a significant effect on primary orthotopic tumor growth was somewhat unexpected. This discrepancy highlights the limitations of *in vitro* models and suggests that the tumor microenvironment of the mammary fat pad provides supportive signals that overcome the growth-inhibitory effects of E-cadherin observed in culture. *In vitro*, E-cadherin re-expression also reduced haptotaxis toward Collagen I, complementing our prior observation that E-cadherin knockout in NST cell lines enhanced Collagen I haptotaxis *In vitro*, E-cadherin re-expression reduced haptotaxis toward Collagen I in ILC cells, a finding that complements our prior observation that E-cadherin knockout in NST cell lines enhanced Collagen I haptotaxis (*32*).

Tail vein injection, a model of systemic dissemination and metastatic outgrowth, yielded both E-cadherin-positive and E-cadherin-negative lesions despite injection of near-universally E-cadherin-positive cells. This implies selective pressure favoring the outgrowth of E-cadherin-negative variants and suggests that E-cadherin acts as a metastasis suppressor in these models. In the MDA-MB-134 cohort, EV-injected mice developed metastases in the reproductive tract, brain, and kidneys, whereas E-cadherin-expressing cells gave rise to fewer metastases. Fewer differences in metastases between EV and E-cadherin-cells were observed in the SUM44 cohort. Although E-cadherin-expressing cells initially showed increased brain tropism, very few of these lesions retained E-cadherin expression again supporting selective pressure for metastatic outgrowth of E-cadherin-negative ILC cells. Overall, these findings indicate that E-cadherin expression reduces metastatic propensity, with more pronounced effects in the MDA-MB-134 model. This is consistent with the marked ULA growth suppression in MDA-MB-134, raising the possibility that E-cadherin-expressing cells undergo anoikis during hematogenous transit, thereby limiting the pool of cells competent to seed distant metastases.

In the MFP model, which more closely recapitulates the clinical sequence of events, E-cadherin restoration strongly decreased metastases to the reproductive tract and GI tract in SUM44 cells, while effects in MDA-MB-134 were less pronounced. As with the tail vein experiments, only a minority of metastases from mice injected with E-cadherin overexpressing cells actually retained E-cadherin expression, and the proportion of E-cadherin-null lesions was substantially greater in the MFP model than in the tail vein cohorts. Despite sorting for both E-cadherin-positive and top-10% RFP-positive cell populations, our sorted E-cadherin-expressing cells gave rise to metastatic lesions that had lost E-cadherin expression. These E-cadherin-negative metastases could have originated from rare E-cadherin-negative cells present in the sorted inoculum, or from E-cadherin-expressing cells that silenced E-cadherin after dissemination. Our experimental design cannot distinguish between these possibilities. The shorter duration of tail vein experiments relative to the MFP/surgical resection model likely accounts for the higher proportion of E-cadherin-null metastases observed in the latter, consistent with prolonged *in vivo* selective pressure against E-cadherin-expressing cells.

The predominant metastatic tropism toward the reproductive tract observed in both MDA-MB-134 and SUM44 models is consistent with the well-documented clinical propensity of ILC to metastasize to the ovaries and peritoneum, a pattern less frequently observed in NST. The selective suppression of reproductive tract metastasis by E-cadherin re-expression may reflect the importance of anoikis resistance for peritoneal dissemination, a metastatic route that requires survival of tumor cells in suspension within the peritoneal cavity. Alternatively, E-cadherin re-expression might alter the homing of ILC cells to the reproductive tract with yet-to-be-discovered mechanism.

Collectively, our data demonstrate that E-cadherin is dispensable for metastasis in ILC, and that its re-expression selectively suppresses metastasis to certain organs. These findings differ from those of Padmanaban et al., who reported E-cadherin as a requirement for metastasis (*23*): the authors showed that E-cadherin loss in NST models reduced cell proliferation, survival, seeding in distant organs, and metastatic outgrowth. In ILC, the reverse holds true: E-cadherin re-expression suppresses growth and metastases. Taken together, our findings and the broader literature underscore that the consequences of E-cadherin loss are highly context-dependent, producing opposing effects on the metastatic process in ILC versus NST.

Given the extensive evidence for E-cadherin’s enabling role in metastasis in epithelial cancers broadly, metastatic recurrence in patients with ILC implies that these tumors have evolved and developed compensatory mechanisms that functionally substitute for E-cadherin-mediated adhesion. Analogous to the approach of Bajrami et al. (*36*) who leveraged E-cadherin CRISPR knockout in NST cells to identify synthetic lethal partners of *CDH1* loss including ROS1, our inducible E-cadherin-overexpression models could be exploited to uncover pathways specifically dependent on E-cadherin loss in ILC.

Our study has several limitations, most notably the absence of analyses examining the effect of E-cadherin expression on bone metastasis. At the time these experiments were conducted, the technical requirements to analyze bone metastasis were not available in our laboratory. In addition, the limited number of models, ie the use of only two ILC cell lines, precludes a deeper understanding of the role of the genetic background in the observed phenotypes. We clearly see a differential response to E-cadherin re-expression between MDA-MB-134 and SUM44 cells which is further complicated by its dependence on the experimental model used. In the tail vein model, which bypasses the primary tumor microenvironment and selectively interrogates survival during hematogenous transit and metastatic seeding, effects were more pronounced in MDA-MB-134 cells, an effect consistent with their greater anoikis sensitivity upon E-cadherin re-expression observed *in vitro*. In contrast, the stronger effects seen in SUM44 cells in the MFP model suggest that E-cadherin re-expression in this line primarily influences earlier steps of the metastatic cascade such as local invasion, intravasation, or interactions with the primary tumor microenvironment, rather than hematogenous survival per se. Collectively, these findings suggest that E-cadherin suppresses metastasis through distinct mechanisms in MDA-MB-134 and SUM44 cells. Future studies should employ additional cell lines models, including syngeneic lines to allow assessment of the immune tumor microenvironment. Such studies should then also include the analysis of bone metastasis. Nevertheless, our study provides new insight into the role of E-cadherin loss in ILC progression, and the inducible models described here represent a valuable resource for identifying novel ILC-specific therapeutic vulnerabilities.

---

**Supplementary Figure S1. E-cadherin and P120-catenin colocalization and immunoblotting reveal 24-hour threshold for adherens junction formation**

A) B) and C) Quantification of immunofluorescence confocal microscopy of E-cadherin staining overlapping with p120-catenin overtime with dox treatment. D) E) and F) Immunoblot using antibodies for E-cadherin, α, β and p120-catenins on whole cell lysate of indicated cells with increasing dox exposure (hours).

**Supplementary Figure S2. Apoptotic response to E-cadherin re-expression is enhanced under ULA conditions**

A) Representative Annexin V and PI FACS staining plots of MDA-MB-134-EV and MDA-MB-134-CDH1 after 4 days in 2D (top row) or ULA (bottom row). B) BCK4 growth curve over 8 days in 2D (line solid) with and without dox (line dashed) in both EV (blue) and CDH1 (red).

**Supplementary Figure S3. E-cadherin does not alter chemotaxis nor invasion phenotypes**

A) Images of crystal violet stained insert with indicated cells from chemotaxis assay toward FBS for 72 hours. B) Quantification of chemotaxis assay (n=3 biological replicates, except for positive control MDA-MB-231 n=2). C) Images of crystal violet stained insert with indicated cells from invasion assay toward Collagen I and ECM for 72 hours. D) Quantification of invasion assays (n=3 biological replicates, except for positive control MM231 n=1) ns, not significant.

**Supplementary Figure S4. Direct contact with Collagen I is required for haptotaxis**

A) Experimental design of contact-dependent haptotaxis assay. B) Images of crystal violet stained insert with SUM44-EV and CDH1 cells from regular transwell with below plate coated with Collagen I. C) Quantification of transwell assay with below plate coated with Collagen I (n=3).

**Supplementary Figure S5. E-cadherin expression inhibits haptotaxis to fibronectin**

A) Experimental design of haptotaxis assay with fibronectin. B) and D) Images of crystal violet stained insert with SUM44-EV and CDH1 cells and other indicated cells from haptotaxis assay toward fibronectin for 72 hours. C) and E) Quantification of haptotaxis assay showing representative data from 1-2 experiments (n=2). P values are from ordinary two-way ANOVA. ns, not significant.

**Supplementary Figure S6. RFP/Luciferase labeling does not alter dox-induced E-cadherin expression or proliferation**

A) Immunoblot for E-cadherin of RFP/Luciferase tagged (hashed) EV in blue CDH1 in red for both SUM44 and MDA-MB-134 cell lines. For comparison their counterpart not tagged CDH1 cell lines are represented in solid red. B) Growth assay in 2D of indicated cell lines with (line dashed) and without (line solid) the RFP/Luciferase tag.

**Supplementary Figure S7. Doxycycline induced E-cadherin expression is maintained *in vivo* without altering Ki67 levels in primary tumors**

A) & B) Immunochemistry for E-cadherin in primary tumor of 3 additional mice from each EV and CDH1 group. C) & D) Immunohistochemistry for Ki67 of EV and CDH1 tumors.

**Supplementary Figure S8. Extracellular matrix alterations through E-cadherin overexpression**

A) & B) Trichrome staining of EV and CDH1 tumors. C) &D) Quantification of trichrome staining showing the % area of collagen. Statistical analysis was performed using unpaired t test. ns, not significant.

**Supplementary Figure S9. IHC of E-cadherin negative CDH1 tumors following TV injection**

A) & B) IHC validation of CDH1 lesion that are E-cadherin negative. CK19 marks human cells to identify the lesions within the tissue.

**Supplementary Figure S10. IHC of E-cadherin negative CDH1 tumors following MFP injection**

A) Table summarizing mice survival pre-surgery, surgery success rate and successful harvest post-surgery. B) & C) IHC validation of CDH1 lesion that are E-cadherin negative. CK19 marks human cells to identify the lesions within the tissue.

## Authors’ Contributions

Conception and design: L. Savariau, N. Tasdemir, S. Oesterreich, A.V. Lee

Development of methodology: L. Savariau, N. Tasdemir, A. Elangovan, D. J. S. John Mary, K. Ding, J. Hooda, J.M. Atkinson, S. Oesterreich, A.V. Lee

Acquisition of data: L. Savariau, A. Elangovan

Analysis and interpretation of data: L. Savariau, A. Elangovan, K. Ding, I. Thale, J.M. Atkinson, S. Oesterreich, A.V. Lee

Writing, review, and/or revision of the manuscript: L. Savariau, I. Thale, N. Tasdemir, A. Elangovan, K. Ding, J.M. Atkinson, S. Oeterreich, A.V. Lee

Administrative, technical, or material support: L. Savariau, A. Elangovan, J.M. Atkinson, S. Oesterreich, A.V. Lee

Study supervision: L.Savariau, S. Oesterreich, A.V. Lee

## Acknowledgements

The authors would like to thank Jian Chen, and Beth Knapick for technical assistance and laboratory support.

## Citation

1. R. L. Siegel, T. B. Kratzer, A. N. Giaquinto, H. Sung, A. Jemal, Cancer statistics, 2025. CA Cancer J Clin 75, 10–45 (2025).

2. A. B. Mariotto, L. Enewold, J. Zhao, C. A. Zeruto, K. R. Yabroff, Medical Care Costs Associated with Cancer Survivorship in the United States. Cancer Epidemiol Biomarkers Prev 29, 1304–1312 (2020).

3. A. Tremont, J. Lu, J. T. Cole, Endocrine Therapy for Early Breast Cancer: Updated Review. Ochsner J 17, 405–411 (2017).

4. V. Guarneri, P. Conte, Metastatic breast cancer: therapeutic options according to molecular subtypes and prior adjuvant therapy. Oncologist 14, 645–656 (2009).

5. R. Barroso-Sousa, O. Metzger-Filho, Differences between invasive lobular and invasive ductal carcinoma of the breast: results and therapeutic implications. Ther Adv Med Oncol 8, 261–266 (2016).

6. W. P. Evans, L. J. Warren Burhenne, L. Laurie, K. F. O’Shaughnessy, R. A. Castellino, Invasive lobular carcinoma of the breast: mammographic characteristics and computer-aided detection. Radiology 225, 182–189 (2002).

7. H. He, A. Gonzalez, E. Robinson, W. T. Yang, Distant metastatic disease manifestations in infiltrating lobular carcinoma of the breast. AJR Am J Roentgenol 202, 1140–1148 (2014).

8. M. Inoue et al., Specific sites of metastases in invasive lobular carcinoma: a retrospective cohort study of metastatic breast cancer. Breast Cancer 24, 667–672 (2017).

9. A. Mathew et al., Distinct Pattern of Metastases in Patients with Invasive Lobular Carcinoma of the Breast. Geburtshilfe Frauenheilkd 77, 660–666 (2017).

10. M. Blohmer et al., Patient treatment and outcome after breast cancer orbital and periorbital metastases: a comprehensive case series including analysis of lobular versus ductal tumor histology. Breast Cancer Res 22, 70 (2020).

11. G. Arpino, V. J. Bardou, G. M. Clark, R. M. Elledge, Infiltrating lobular carcinoma of the breast: tumor characteristics and clinical outcome. Breast Cancer Res 6, R149–156 (2004).

12. N. Biglia et al., Increased incidence of lobular breast cancer in women treated with hormone replacement therapy: implications for diagnosis, surgical and medical treatment. Endocr Relat Cancer 14, 549–567 (2007).

13. N. Wasif, M. A. Maggard, C. Y. Ko, A. E. Giuliano, Invasive lobular vs. ductal breast cancer: a stage-matched comparison of outcomes. Ann Surg Oncol 17, 1862–1869 (2010).

14. J. Timbres et al., Survival Outcomes in Invasive Lobular Carcinoma Compared to Oestrogen Receptor-Positive Invasive Ductal Carcinoma. Cancers (Basel) 13, (2021).

15. S. Oesterreich et al., Clinicopathological Features and Outcomes Comparing Patients With Invasive Ductal and Lobular Breast Cancer. J Natl Cancer Inst 114, 1511–1522 (2022).

16. B. C. Pestalozzi et al., Distinct clinical and prognostic features of infiltrating lobular carcinoma of the breast: combined results of 15 International Breast Cancer Study Group clinical trials. J Clin Oncol 26, 3006–3014 (2008).

17. G. Ciriello et al., Comprehensive Molecular Portraits of Invasive Lobular Breast Cancer. Cell 163, 506–519 (2015).

18. C. Desmedt et al., Genomic Characterization of Primary Invasive Lobular Breast Cancer. J Clin Oncol 34, 1872–1881 (2016).

19. N. Cancer Genome Atlas, Comprehensive molecular portraits of human breast tumours. Nature 490, 61–70 (2012).

20. H. Dopeso et al., Genomic and epigenomic basis of breast invasive lobular carcinomas lacking CDH1 genetic alterations. NPJ Precis Oncol 8, 33 (2024).

21. J. H. Venhuizen, F. J. C. Jacobs, P. N. Span, M. M. Zegers, P120 and E-cadherin: Double-edged swords in tumor metastasis. Semin Cancer Biol 60, 107–120 (2020).

22. Z. Li, S. Yin, L. Zhang, W. Liu, B. Chen, Prognostic value of reduced E-cadherin expression in breast cancer: a meta-analysis. Oncotarget 8, 16445–16455 (2017).

23. V. Padmanaban et al., E-cadherin is required for metastasis in multiple models of breast cancer. Nature 573, 439–444 (2019).

24. A. E. McCart Reed et al., An epithelial to mesenchymal transition programme does not usually drive the phenotype of invasive lobular carcinomas. J Pathol 238, 489–494 (2016).

25. P. Jambal et al., Estrogen switches pure mucinous breast cancer to invasive lobular carcinoma with mucinous features. Breast Cancer Res Treat 137, 431–448 (2013).

26. M. J. Sikora et al., Invasive lobular carcinoma cell lines are characterized by unique estrogen-mediated gene expression patterns and altered tamoxifen response. Cancer Res 74, 1463–1474 (2014).

27. N. Tasdemir et al., Proteomic and transcriptomic profiling identifies mediators of anchorage-independent growth and roles of inhibitor of differentiation proteins in invasive lobular carcinoma. Sci Rep 10, 11487 (2020).

28. N. Tasdemir et al., Comprehensive Phenotypic Characterization of Human Invasive Lobular Carcinoma Cell Lines in 2D and 3D Cultures. Cancer Res 78, 6209–6222 (2018).

29. K. Stock et al., Capturing tumor complexity in vitro: Comparative analysis of 2D and 3D tumor models for drug discovery. Sci Rep 6, 28951 (2016).

30. P. W. Derksen et al., Somatic inactivation of E-cadherin and p53 in mice leads to metastatic lobular mammary carcinoma through induction of anoikis resistance and angiogenesis. Cancer Cell 10, 437–449 (2006).

31. R. C. Schackmann et al., Cytosolic p120-catenin regulates growth of metastatic lobular carcinoma through Rock1-mediated anoikis resistance. J Clin Invest 121, 3176–3188 (2011).

32. A. Elangovan et al., Loss of E-cadherin Induces IGF1R Activation and Reveals a Targetable Pathway in Invasive Lobular Breast Carcinoma. Mol Cancer Res 20, 1405–1419 (2022).

33. N. Tasdemir et al., Estrogen receptor-positive cell line xenograft models recapitulate metastatic dissemination and endocrine response of invasive lobular breast carcinoma. bioRxiv, (2026).

34. T. T. Onder et al., Loss of E-cadherin promotes metastasis via multiple downstream transcriptional pathways. Cancer Res 68, 3645–3654 (2008).

35. R. A. van de Ven et al., Nuclear p120-catenin regulates the anoikis resistance of mouse lobular breast cancer cells through Kaiso-dependent Wnt11 expression. Dis Model Mech 8, 373–384 (2015).

36. I. Bajrami et al., E-Cadherin/ROS1 Inhibitor Synthetic Lethality in Breast Cancer. Cancer Discov 8, 498–515 (2018).

