## Supplemental Figures for "Restoration of E-cadherin Expression Alters Metastatic Organotropism in Invasive Lobular Breast Carcinoma Models"

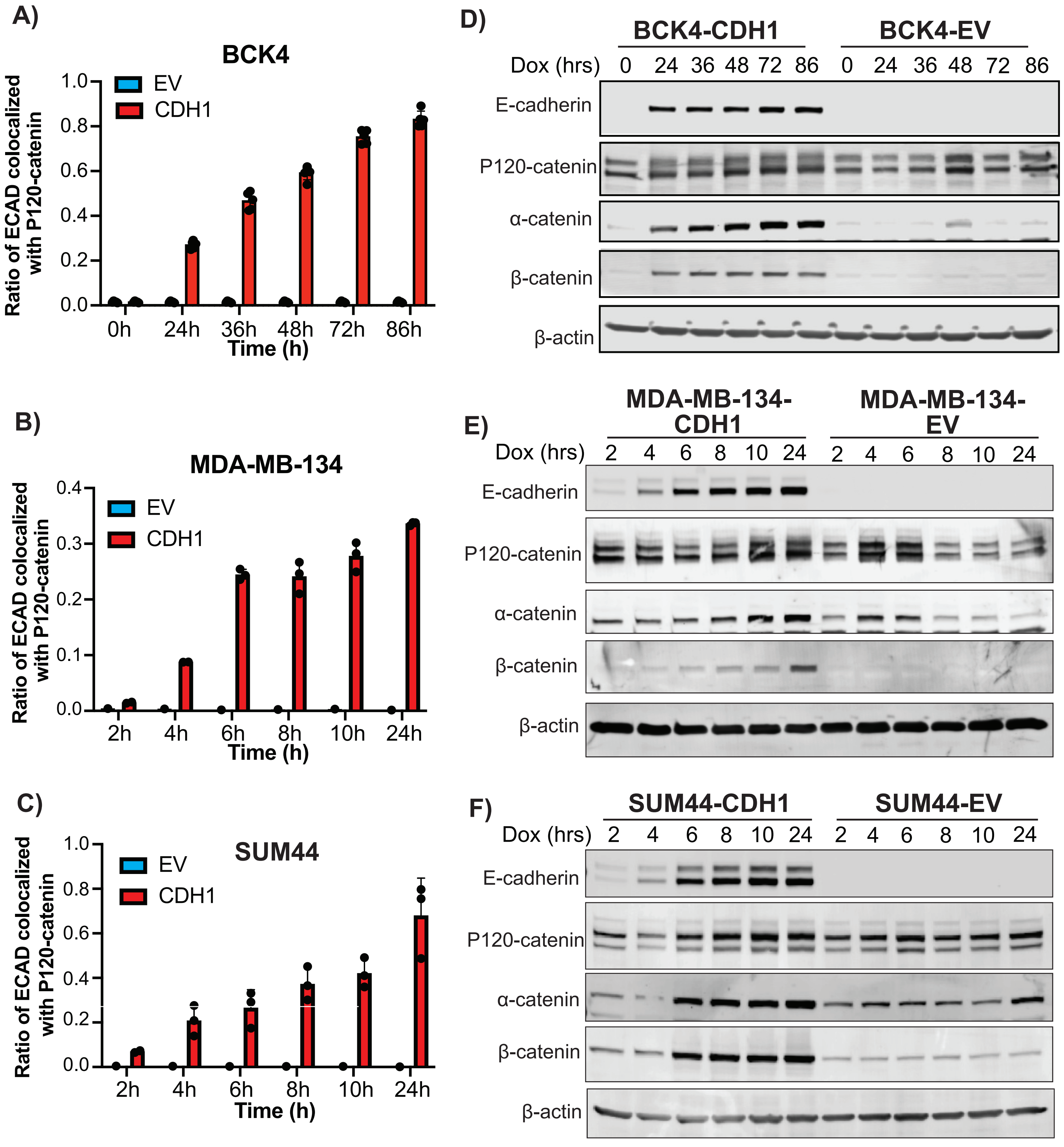


**Supplementary Figure S1. E-cadherin and P120-catenin colocalization and immunoblotting reveal 24-hour threshold for adherens junction formation**

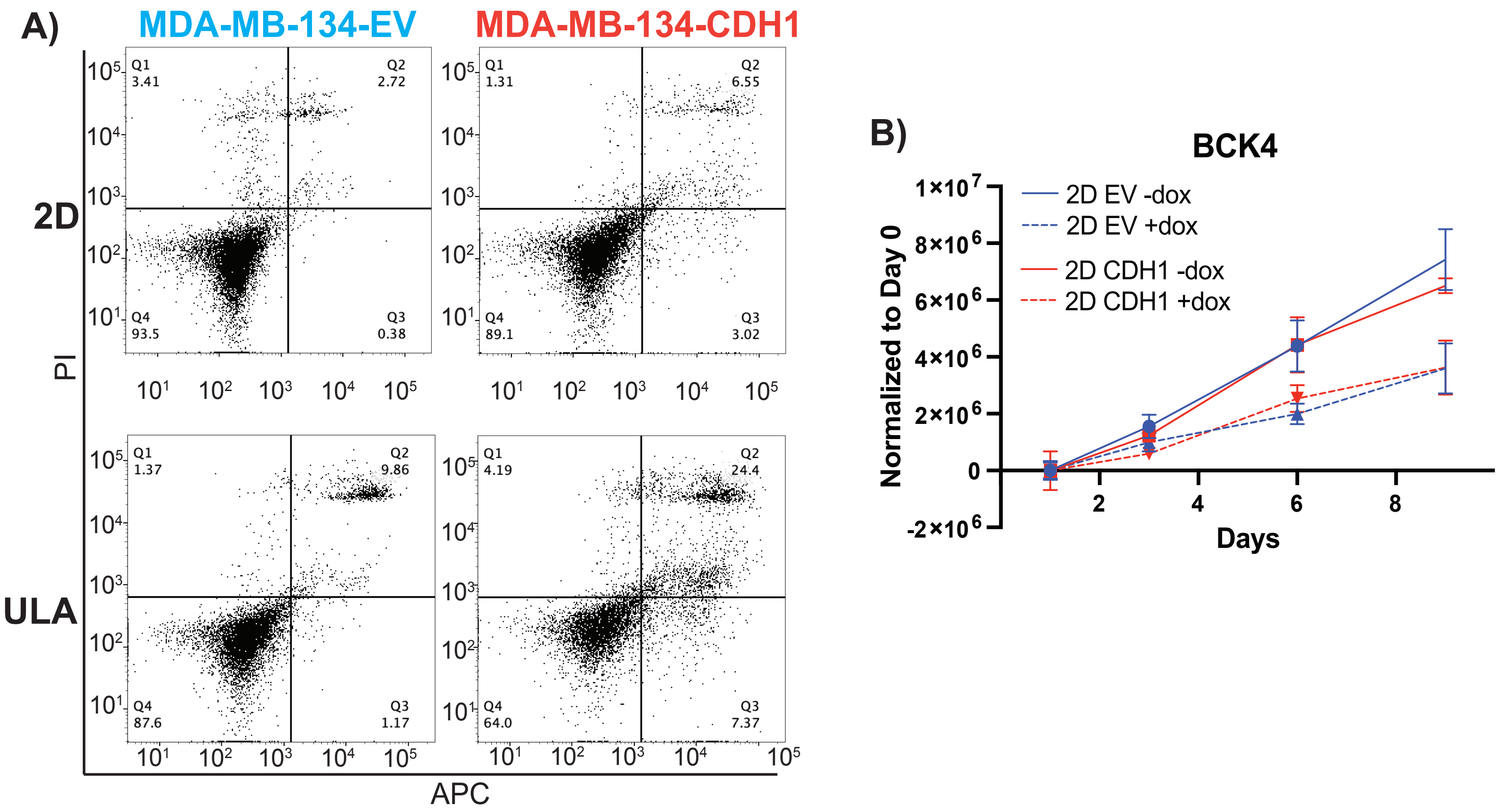


**Supplementary Figure S2. Apoptotic response to E-cadherin re-expression is enhanced under ULA conditions**

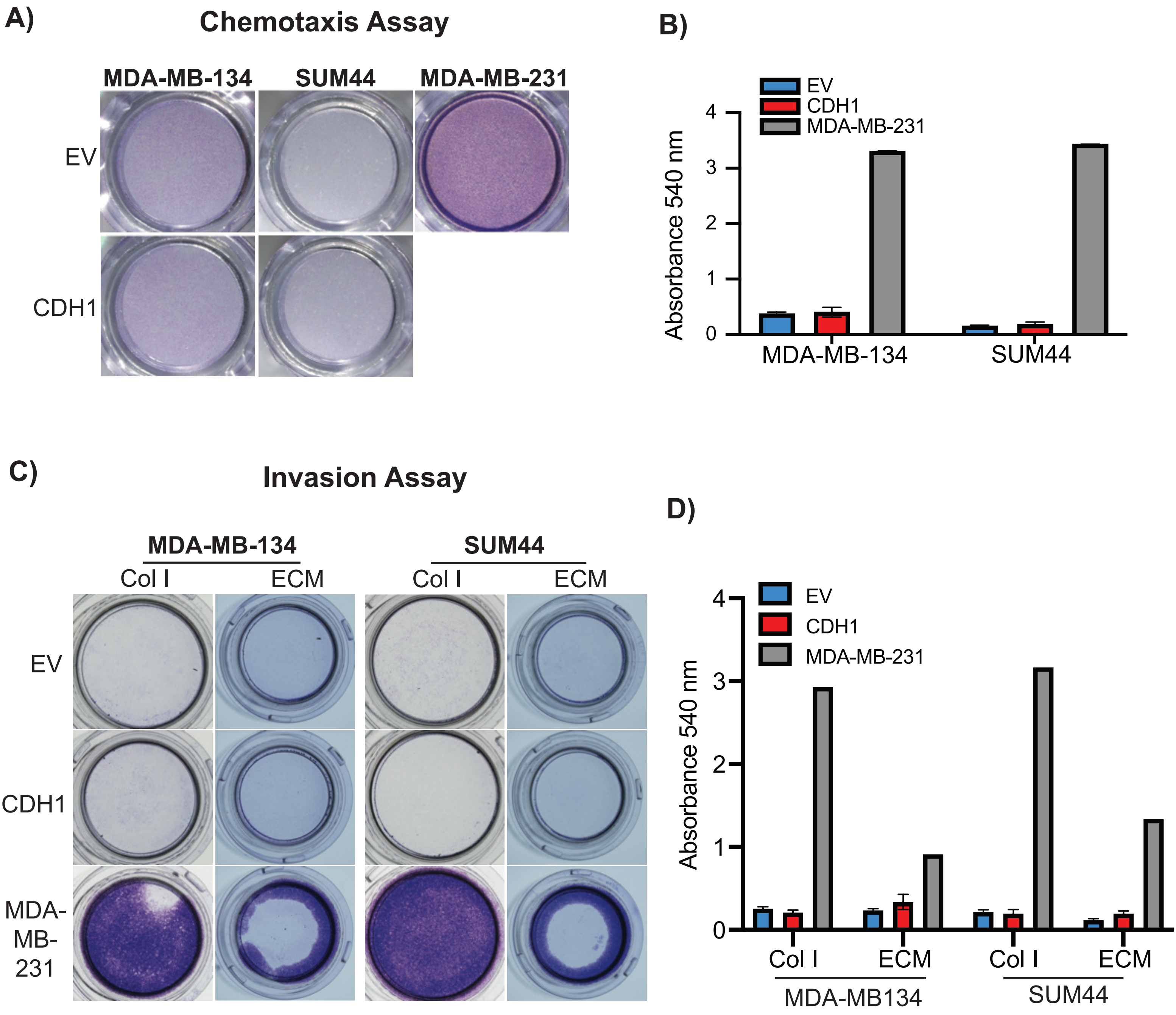


**Supplementary Figure S3. E-cadherin does not alter chemotaxis nor invasion phenotypes**

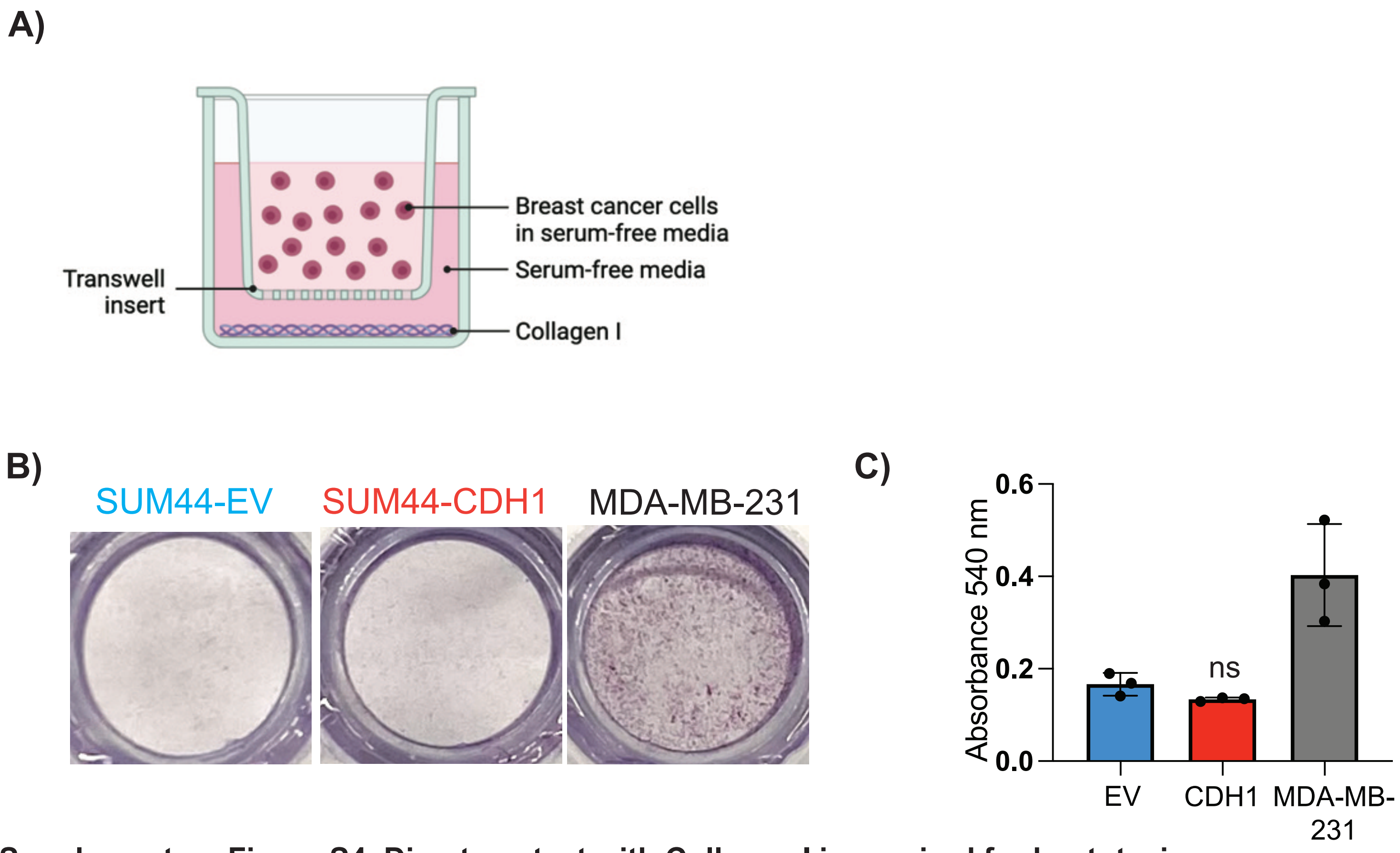


**Supplementary Figure S4. Direct contact with Collagen I is required for haptotaxis**

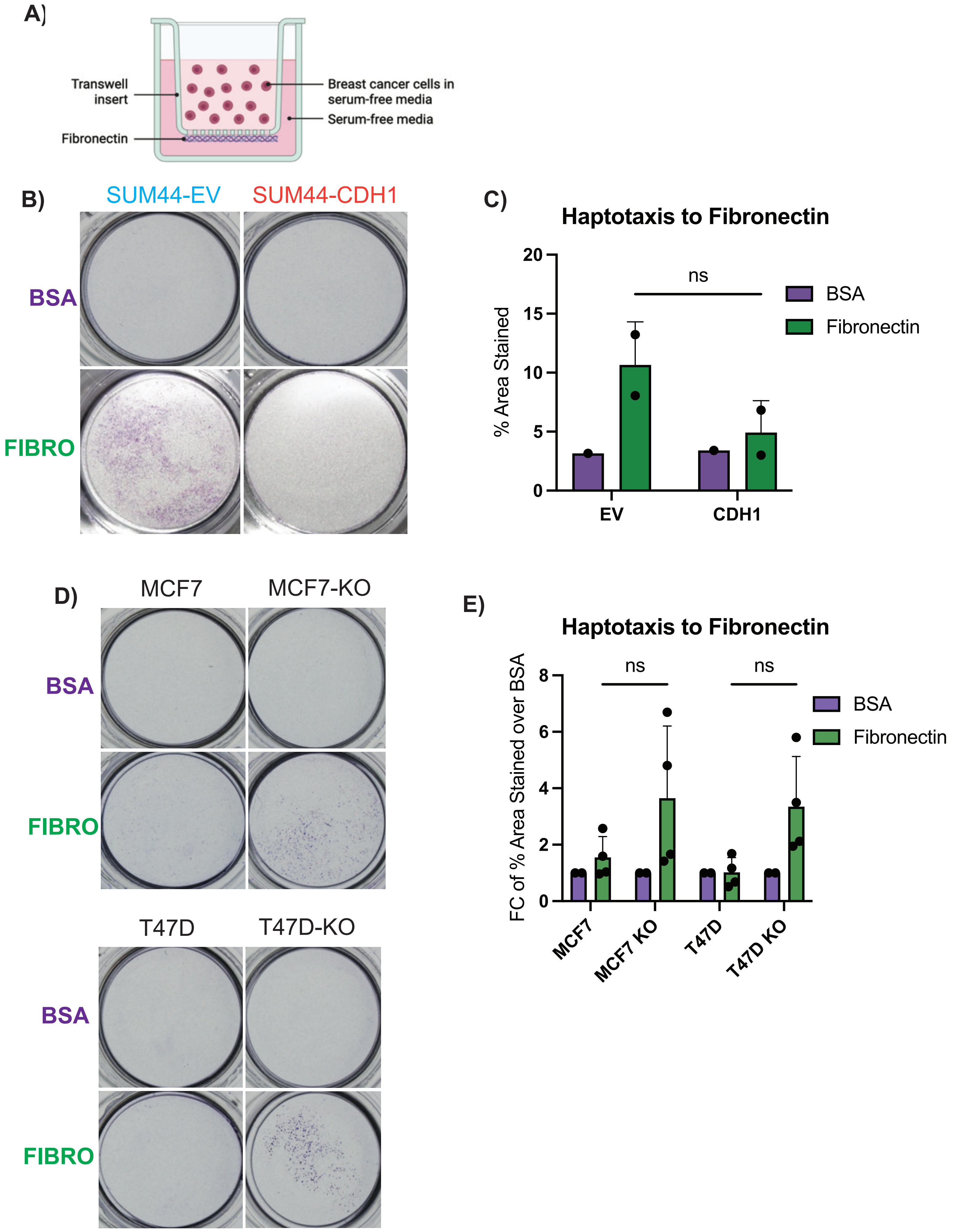


**Supplementary Figure S5. E-cadherin expression inhibits haptotaxis to fibronectin**

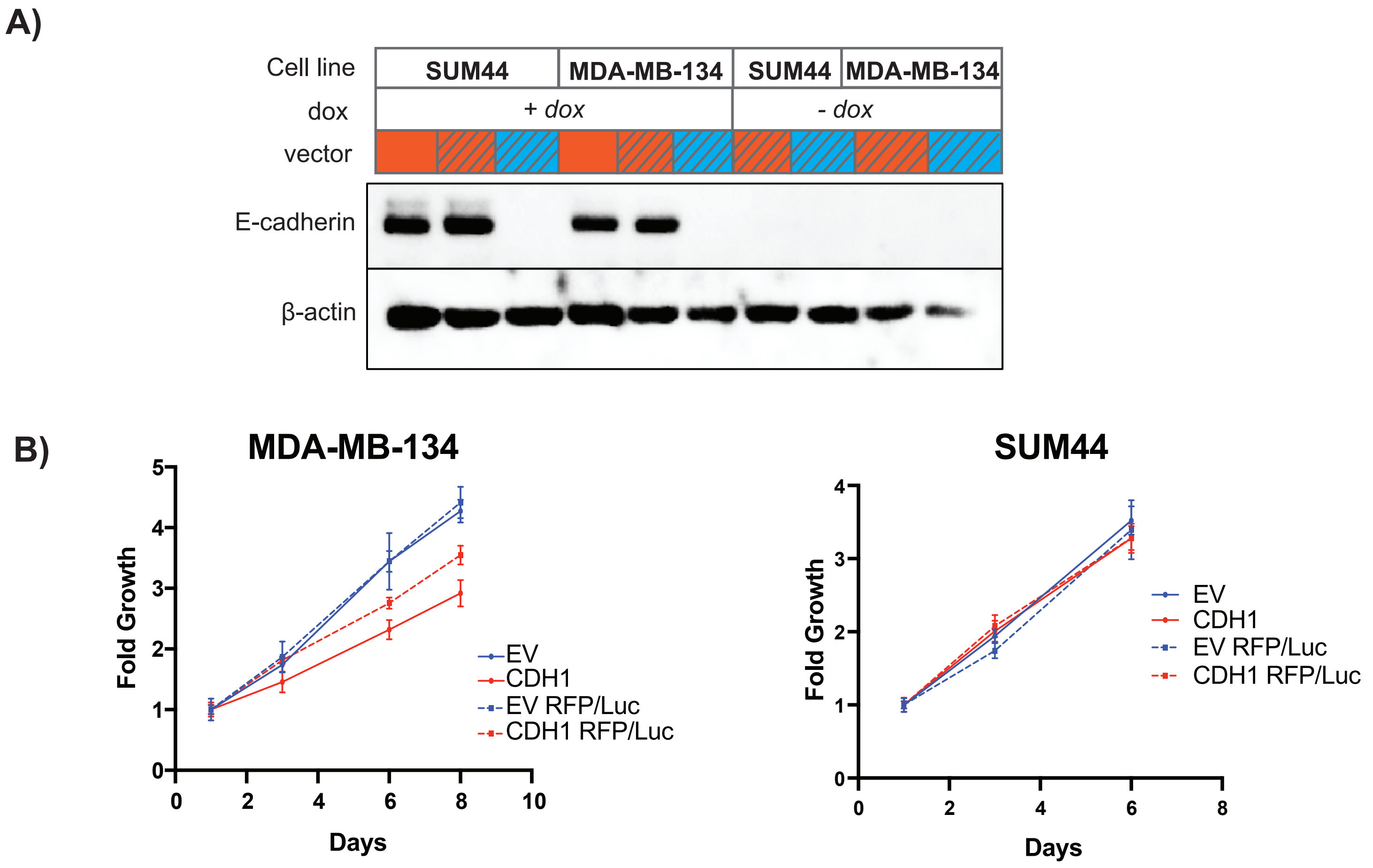


**Supplementary Figure S6. RFP/Luciferase labeling does not alter dox-induced E-cadherin expression or proliferation**

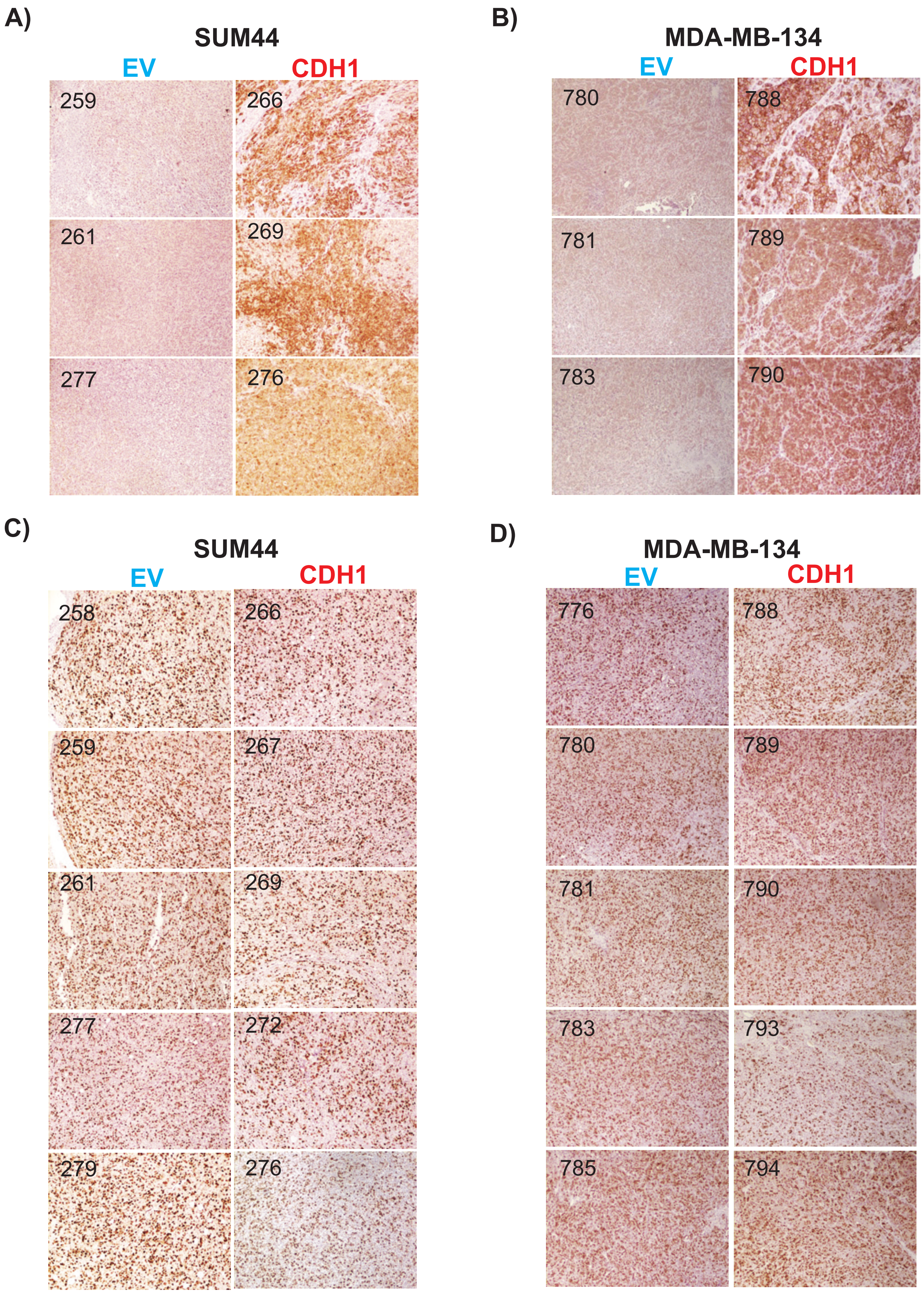


**Supplementary Figure S7. Doxycycline induced E-cadherin expression is maintained *in vivo* without altering Ki67 levels in primary tumors**

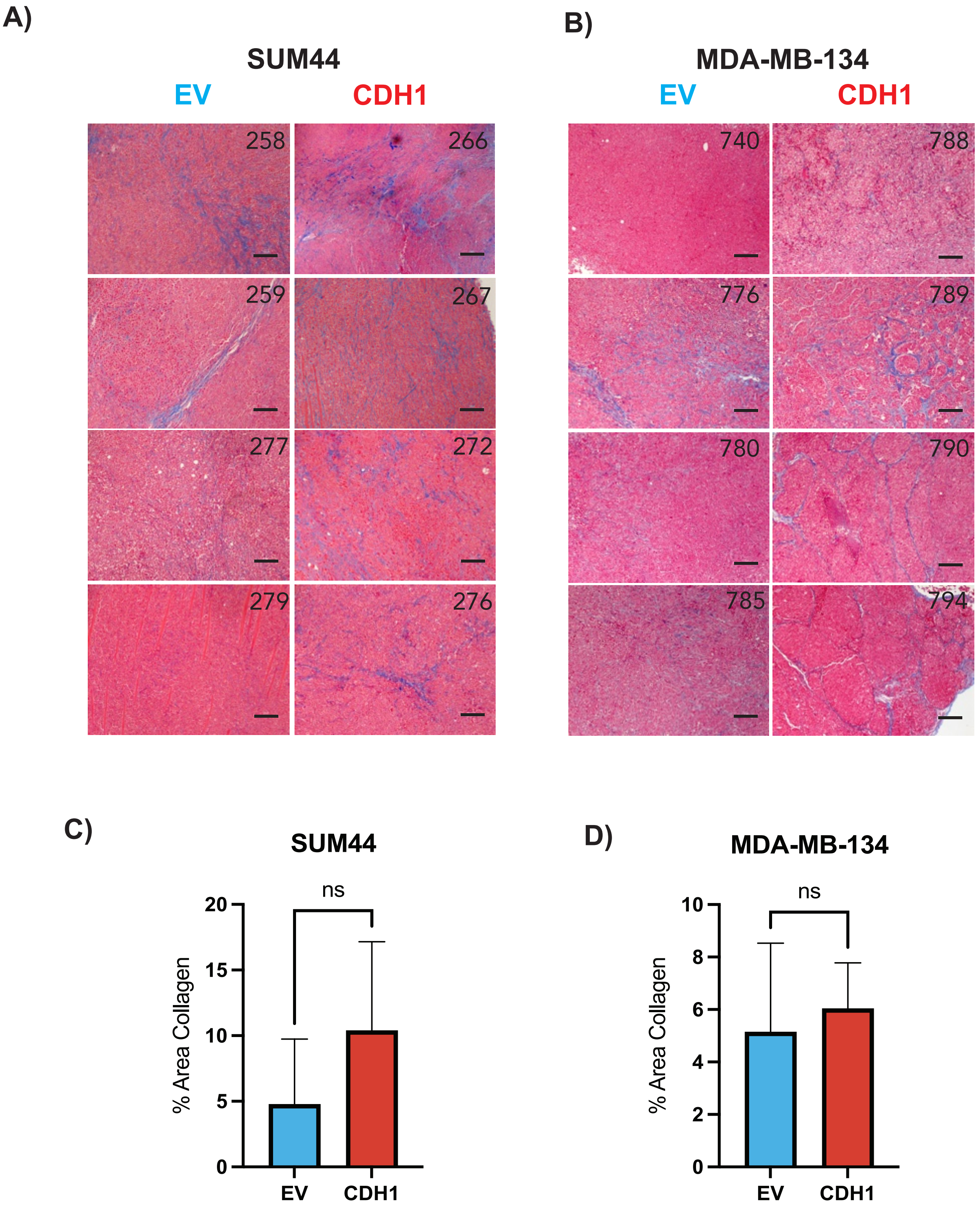


**Supplementary Figure S8. Extracellular matrix alterations through E-cadherin overexpression**

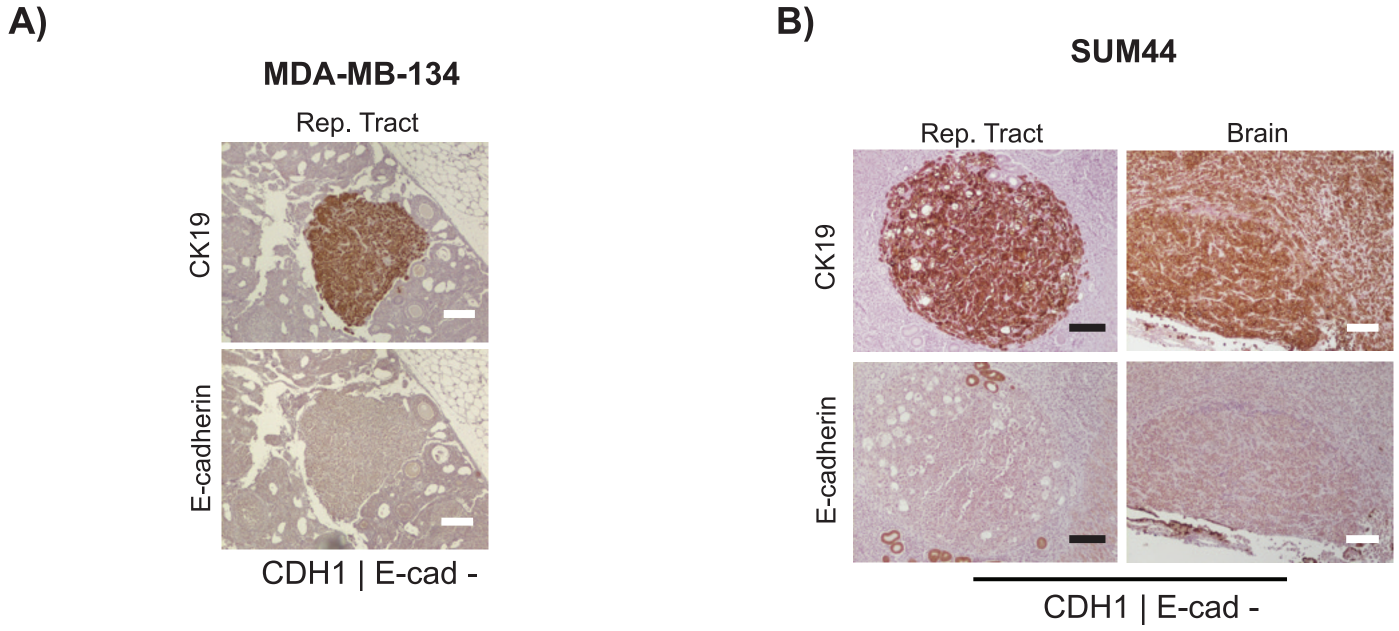


**Supplementary Figure S9. IHC of E-cadherin negative CDH1 tumors following TV injection**

A)

| | SUM44 | MDA-MB-134 |
|---|---|---|
| ***Mice survival pre-surgery*** | 96% | 100% |
| ***Surgery success rate*** | 87% | 91.7% |
| ***Successful harvest post-surgery*** | 62% | 79% |

B)

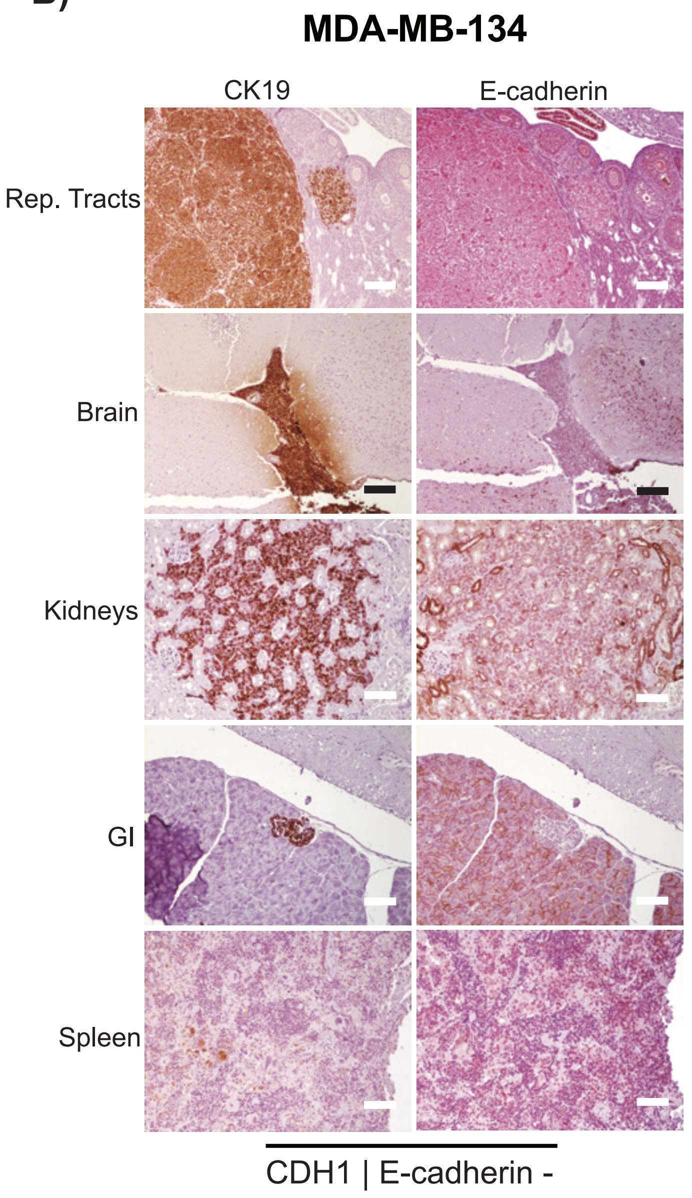


C)

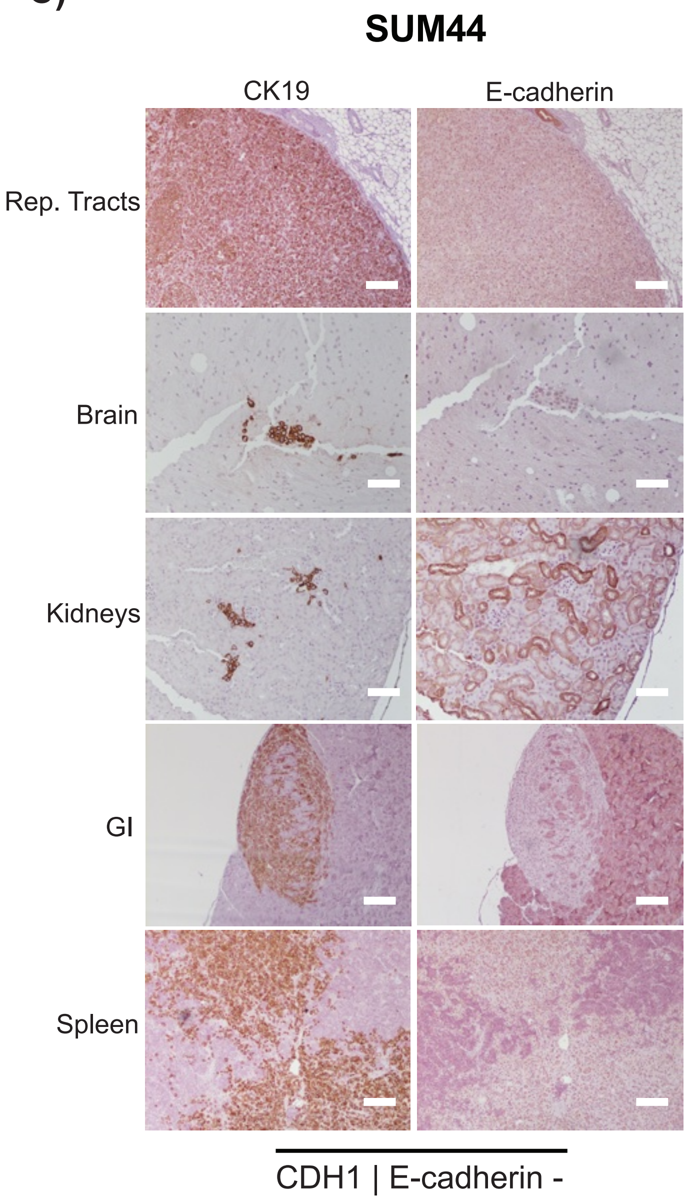


**Supplementary Figure S10. IHC of E-cadherin negative CDH1 tumors following MFP injection**
